# Conditional, tissue-specific CRISPR/Cas9 vector system in zebrafish reveal the role of neuropilin-1b in heart regeneration

**DOI:** 10.1101/2023.01.16.524222

**Authors:** Ramcharan Singh Angom, Ying Wang, Enfeng Wang, Shamit Dutta, Debabrata Mukhopadhyay

## Abstract

CRISPR/Cas9 technology-mediated genome editing has significantly improved the targeted inactivation of genes *in vitro* and *in vivo* in many organisms. In this study, we have reported a novel CRISPR-based vector system for conditional tissue-specific gene ablation in zebrafish. Specifically, the cardiac-specific *cardiac myosins light chain 2 (cmlc2*) promoter drives Cas9 expression to silence the *neuropilin-1(nrp1*) gene in cardiomyocytes in a heat-shock inducible manner. This vector system establishes a unique tool to regulate the gene knockout in both the developmental and adult stages and hence, widens the possibility of loss-of-function studies in zebrafish at different stages of development and adulthood. Using this approach, we investigated the role of neuropilin isoforms *nrp1a* and *nrp1b* in response to cardiac injury and regeneration in adult zebrafish hearts. We observed that both the isoforms (*nrp1a* and *nrp1b*) are upregulated after the cryoinjury. Interestingly, the *nrp1b*-knockout significantly altered heart regeneration and impaired cardiac function in the adult zebrafish, demonstrated by reduced heart rate (ECG), ejection fractions, and fractional shortening. In addition, we show that the knockdown of *nrp1b* but not *nrp1a* induces activation of the cardiac remodeling genes in response to cryoinjury. To our knowledge, this is the first study where we have reported a heat shock-mediated conditional knockdown of *nrp1a* and *nrp1b* isoforms using CRISPR/Cas9 technology in the cardiomyocyte in zebrafish, and furthermore have identified a crucial role for *nrp1b* isoform in zebrafish cardiac remodeling and eventually heart function in response to injury.

## Introduction

CRISPR/Cas9 technology has become a powerful tool for targeted genome editing^1^; adapted from the bacterial immune mechanism^2^, it utilizes the advantage of the RNA-dependent detection of specific DNA sequences by Cas9 endonuclease. The designed guide RNA (gRNA) comprises a 3’ secondary structure that can interact with Cas9. It also contains a 20 bases long 5’ seed sequence that leads to targeting. The Cas9 screens the genome and cleaves it within sequences complementary to the seed sequence, which are immediately followed by the adjacent protospacer motif (PAM) NGG^3^ Previously, genes can efficiently be inactivated by transfection of gRNAs and Cas9 mRNA or DNA into bacteria, human or mouse cells^4–6^.

Zebrafish (Danio rerio) has emerged as a popular as a model system for various biological studies, including cardiovascular diseases^7–9^. Specific characteristics such as transparent embryos, a short life cycle^10^, a highly conserved innate immune system with humans^11^, and ease of genetic manipulation^12^ make the zebrafish ideal for live imaging and dissecting new mechanisms regulating cardiovascular development. The application of the CRISPR/Cas9 tool in zebrafish is well documented. Previously, injection of gRNAs and Cas9 mRNA into one-cell stage embryos has been shown to produce indels at the target loci with comparatively high, although varying, rate of occurrence ^13,14^. CRISPR-mediated mutations in zebrafish are heritable, allowing mutant strain’s speedy generation. Since gene deactivation by CRISPR/Cas9 tool is comprehensive and stable, this technology has provided an effective complementary method to morpholinos for loss-of-function studies in zebrafish, especially at subsequent stages of development. As the loss of some genes leads to embryonic lethality, there is a strong requirement to generate conditional, tissue-specific knockouts in the field. Previously, Ablain et al. reported a CRISPR-based vector system that enables stable, tissue-specific gene inactivation *in vivo^15^*. In this study, we report a novel CRISPR-based vector system (modified from the previous vector described by Ablain et al. ^15^ for heat-inducible, cardiomyocyte-specific gene ablation in adult zebrafish. In this vector, the cardiac-specific, *cardiac myosin light chain 2 (cmlc2*) promoter drives the *Cas9* expression to silence the *neuropilin-1(nrp1*) gene in cardiomyocytes in a heat-shock inducible manner. This vector system establishes a unique tool to knock out genes in both early and adult stages, expanding the likelihood of loss-of-function studies in zebrafish at different stages of development.

In contrast to mammals, adult zebrafish hearts can regenerate after injury. Neuropilins are co-receptors that have been reported previously to play crucial roles in zebrafish heart regeneration^16^, heart failure in mice^17^, and electrical remodeling after myocardial infarction in rats^18^. But the cell-specific functions of nrp1 have not been described. In this study, we have investigated the role of neuropilin isoforms, including *nrp1a* and *nrp1b* in cardiomyocytes during cardiac injury and regeneration in adult zebrafish hearts. Our results showed that both the isoforms (*nrp1a* and *nrp1b*) are upregulated in the adult zebrafish heart after the cryoinjury. Cardiomyocyte specific *nrp1b* is required explicitly for cardiac repair in adult zebrafish.

In this study, we employed the novel, conditional CRISPR/Cas9 mediated tissue-specific gene inactivation system and showed that *nrp1b* isoform played a crucial role in adult zebrafish heart repair and function after injury. To our knowledge, this is the first study that described the heat shock inducible, conditional and spatiotemporal knockout of tissue-specific genes in zebrafish. This tool can help study tissue-specific gene inactivation in a spatiotemporal manner.

## Method

### Zebrafish Maintenance

Wildtype zebrafish (*Danio rerio*) were maintained at the zebrafish core facility at Mayo Clinic, Jacksonville, FL. The fish room was held on light control of 14 h of light in a day at 29 °C. Adult zebrafish were kept in 2.75 L and 6 L tanks on a water recirculation rack system with a male-to-female ratio of 1:2. Adult zebrafish were fed a range of dry fish food. Adult male and female zebrafish were separated for one week before breeding. Embryos were harvested by breeding two females and one female. The animal experiments were performed as per the Mayo Clinic-approved IACUC protocol number A000018914-14.

### Construction of a heat shock inducible tissue-specific CRISPR vector for gene targeting in zebrafish

We derived the current vector from^15^. Briefly, we PCR amplified and cloned the zebrafish Hsp70 promoter from the AB zebrafish wild type strain cDNA followed by a NheI site into the pcmlc2:GFP destination vector^15,19^ using ClaI and KpnI enzymes. The gRNA scaffold used by the Joung lab^14^ was modified to replace BsaI enzyme sites with sites for BseRI, which was inserted at the start site of the U6 promoter as described earlier^15^. The 20-bp target sequence of interest was designed using the online tool CHOPCHOP (http://chopchop.cbu.uib.no/). The sgRNA was cloned into the gRNA scaffold. We combined this with the middle-entry vector containing the Cas9 (codon-optimized for zebrafish) developed by the Chen lab^20^. The Cas9 can be introduced under any promoter of interest into pcmlc2:GFP and U6:gRNA in a single cloning step by using multisite Gateway cloning technology^21^.

### Heat shock treatment

Heat shock was performed by transferring the zebrafish (Both groups) from 28.5°C to 37°C embryo water and incubating at 37°C for 30 minutes. Briefly, to induce the heat shock promoter expression, heat treatment was given by raising the water temperature from 28.5°C to 38°C over 15 min, then maintaining at 38°C for 30 min, and finally decreasing it to 28.5°C over the next 15 min in a programmable water bath as described^22,23^. The heat shock treatment administered to Cas9 transgenic embryos induced the gRNA expression.

### mRNA and gRNA synthesis

Cas9 mRNA was produced by in vitro transcription from a pCS2 Cas9 vector as described previously^20^ using mMESSAGE mMACHINE SP6 kit (Thermofisher Scientific). The gRNAs target sequence was identified by using the online tool CHOPCHOP^24^, and was produced following established methods^14^ Three target sequences were tested for each isoform. The Hsp70 and Cmlc2 promoters were PCR amplified from the cDNA established from wild type AB embryos using the following primers: Hsp70: 5’-TCAGGGGTGTCGCTTGGTTATTTCC-3’ and 5’-ATTTATACAATTTATGGTGCAATTG-3’, cmlc2: 5’-GCTTAAATCAGTTGTGTTAAATAAGAGAC-3’, and 5’-GGTCACTGTCTTGCTTTGCTG-3’ and The cDNA sequence was then cloned into a middle entry clone for Gateway and inserted into the pDest Tol2 pA2 vector between an SP6 promoter and a polyA sequence as described earlier^15^

### Microinjections

20 pg of CRISPR DNA constructs and 20 pg of Tol2 mRNA were injected into one-cell stage wildtype zebrafish embryos (AB strain) by using the Harvard apparatus’s picoinjector (PLI-100). Post microinjection, embryos were raised in the embryo water at 28°C-29°C till adulthood.

Each component of the CRISPR construct was confirmed by sanger sequencing and restriction digestion before final assembly, as described in Ablain et al., 2015^15^. The CRISPR editing was confirmed by sanger sequencing and T7E1 mutagenesis assay. CRISPR/Cas9 mediated nrp1a and nrp1b ablation was confirmed by Western CRISPR/cas9 validation

### T7E1 Mutagenesis Assay

T7E1 assay was performed as reported^15^. Briefly, genomic DNA was extracted from 2-day-old embryos using the HotSHOT method. A fragment of approximately 400 bp was amplified from genomic DNA using the following primers. For nrp1a, TGTTTGTTTTTGGTTTGTCAGC and CTTGTGTCGTACTCTATGCAACG For nrp1b: CAGCACACAAGGGTTACCATAA and GTCAATGCATCACTATGGCAGTAACCC.

### Whole-mount in situ hybridization and benzidine staining

We cloned approximately 460 bp long fragments from the 3’ end of the *nrp1a* and *nrp1b* gene into the PCR-TOPO 4 vector (Invitrogen) using primers (*nrp1a* Forward: 5’-GCGCTCTTTCGGTTTCCTTC-3’ *nrp1a* Reverse: 5’-TGAAGAGCTGGTTTCCCGAC-3’), (*nrp1b* forward: 5’-ATCCCAACAACCTGGAGTGC-3’ and *nrp1b* reverse: 5’-GGCGTCCAGCCATTTTCAAG-3’) and synthesized both the sense and anti-sense RNA probes by in vitro transcription. In situ hybridization was performed following established protocols^25^.

### Imaging

Embryos were mounted in 0.8% low melting point agarose containing tricaine (160 mg/L) for imaging. The images were acquired using an Olympus dissecting microscope, Zeiss Axio vision microscope, Raman microscope, and LSM 880 Confocal microscope.

The in situ images in Figure S6 were processed by Adobe photoshop to increase the resolution.

### Cryoinjury

Zebrafish aged 10-12 months were anesthetized in 0.1% tricaine (Sigma Aldrich) and placed in a damp sponge on the ventral side. As described previously, a small incision was made through the chest with iridectomy scissors to access the heart^26^. The ventricular wall was directly frozen by a stainless steel cryoprobe precooled in liquid nitrogen and applied for 23-25 seconds. The tip of the cryoprobe was approximately 6 mm long with a diameter of 0.8 mm.. In order to avoid the complete freezing of the heart by the probe, fish water at room temperature was added to the head of the cryoprobe. Post cryo-injury, the zebrafish were transferred to the normal fish water to recover and were then allowed to regenerate for various times at water temperatures 28°C-29°C.

### ECG recordings and acquisition

All the ECGs were recorded in triplicates before the injury and at the subsequent time points after surgery. To reduce the biological variations, the same animals were used for recordings before the injury and at the following time points after surgery. The zebrafish were anesthetized by immersion in 0.015% tricaine solution for 60 seconds. Anesthetized zebrafish were placed ventral side up in a sponge. Each ECG was recorded for 60-120 seconds using iWORX ECG instrument (iWorx.Inc) by following the previous method^27^, the zebrafish were allowed to recover in water free of tricaine.

### Echocardiogram

Echocardiography was performed on conscious zebrafish using a Vevo 2100 and 3100 high-resolution imaging system with an MS-550S transducer (VisualSonics). Zebrafish were sedated in 0.015% tricaine and immobilized on a sponge immersed in fish water. Imaging was performed on the short axis (perpendicular to the fish) and the long axis (parallel to the fish) using B-mode using the Vevo 2100 and 3100 Cardiac Package, taking the average measurement of three consecutive contractions as described earlier^28^.

### RNA extraction

Zebrafish heart ventricles were dissected and quickly frozen in liquid nitrogen. This was followed by thorough homogenization in 700 ml QIAzol reagent (Invitrogen, Carlsbad, USA) using a motor cordless homogenizer (Kimble, 749540-0000, USA) at maximum speed. Chloroform (250 μl) was added to the homogenized embryo/tissue, followed by shaking vigorously for 15 s and incubating at room temperature for 5 min. The samples were then centrifuged at 13,000 rpm for 10 min at 4°C. The upper aqueous phase containing RNA was carefully transferred to a new tube without disturbing the interface. RNA was precipitated by the addition of an equal volume of 70% ethanol and loaded onto a spin column from an RNeasy mini kit (Qiagen, Valencia, USA) according to the manufacturer’s instructions.

### First-strand cDNA synthesis and RT PCR

Total RNA (1 μg) was reverse transcribed to produce cDNA using a superscript cDNA Synthesis kit (Biorad) as described in the manual. In all cases, the qRT-PCRs were performed using SYBR green. Standard reactions (10μl) were assembled as follows: 5 μl of SYBR green qPCR supermix-with Rox (Invitrogen), 1 μl of forward primer (5 μM), 1μl of reverse primer (5 μM), 1 μl of the template and 2.0 μl nuclease-free water. Templates were 1:10 diluted cDNA samples, and in the case of negative controls, cDNAs were replaced by nuclease-free water. All real-time assays were carried out in triplicate using an Applied Biosystems QuantStudio real-time PCR platform using the primers listed in Table 1. Forty amplification cycles were performed, with each cycle consisting of 94 °C for 15 s followed by 60 °C for 1 min. The primers used are listed in Supplemental table 1.

### Histology

For Hematoxylin and Eosin staining, the heart tissue was fixed in 4% PFA overnight at 4°C and submitted to the core facility at Mayo Clinic, Jacksonville. Briefly, the tissues were dehydrated and embedded in paraffin blocks. Sections were cut at the thickness of 5 μm, rehydrated, and stained with Mayer’s Hamatoxyline for 12 minutes. The nuclear staining was differentiated for 5 seconds in 0.37% HCl prepared in 70% ethanol, and the slides were washed in running tap water for 10 minutes. The protein staining was obtained by incubating for 10 minutes in 0.1% Eosin Y solution in water with a drop of acidic acid, followed by a rapid wash in water. The tissue sections were dehydrated in a water/ethanol series, cleared in xylol, and mounted in Entelan medium (Merck). For morphometric analysis, all consecutive sections of 5 hearts per group were photographed using an Aperio scanner (Leica). The infarct region and the intact myocardium were demarcated, and the sizes were measured using ImageJ software. The percentage of the infarct size relative to the entire ventricle was calculated.

### Statistics

All analyses were performed using GraphPad Prism 9.0 (GraphPad Software Inc., La Jolla, CA, USA). Experiments were routinely repeated at least three times. All values are expressed as the mean ± SD. Statistical differences were determined to be significant at p < 0.05. For comparison between the two groups, a paired or unpaired two-tailed Student’s *t*-test was performed. For the heart rate analysis in Figure 4, statistical significance was evaluated with Prism 9.0 software by using Two-way ANOVA (multiple comparisons).

## Results

### nrp1a and nrp1b are expressed in the heart of zebrafish embryos and adult zebrafish

We first checked *nrp1a* and *nrp1b* mRNA levels in the developing zebrafish hearts using *in situ* hybridization. Both nrp1a and nrp1b were observed to be expressed in the brain zebrafish embryo at 24h and 48h (Fig 1A). The nrp1b isoform was expressed in the heart of 48 hpf and 72 hpf embryos (Fig 1A). The nrp1a isoform also showed expression in the hearts at 72 hpf (Fig 1A). As shown in Fig 1B, the immunostaining shows that nrp1 isoforms, including *nrp1a* and *nrp1b* were ubiquitously expressed in the ventricles and the trabecular muscles in the cardiac atrium and ventricle of 12-month-old zebrafish.

**Figure 1.**
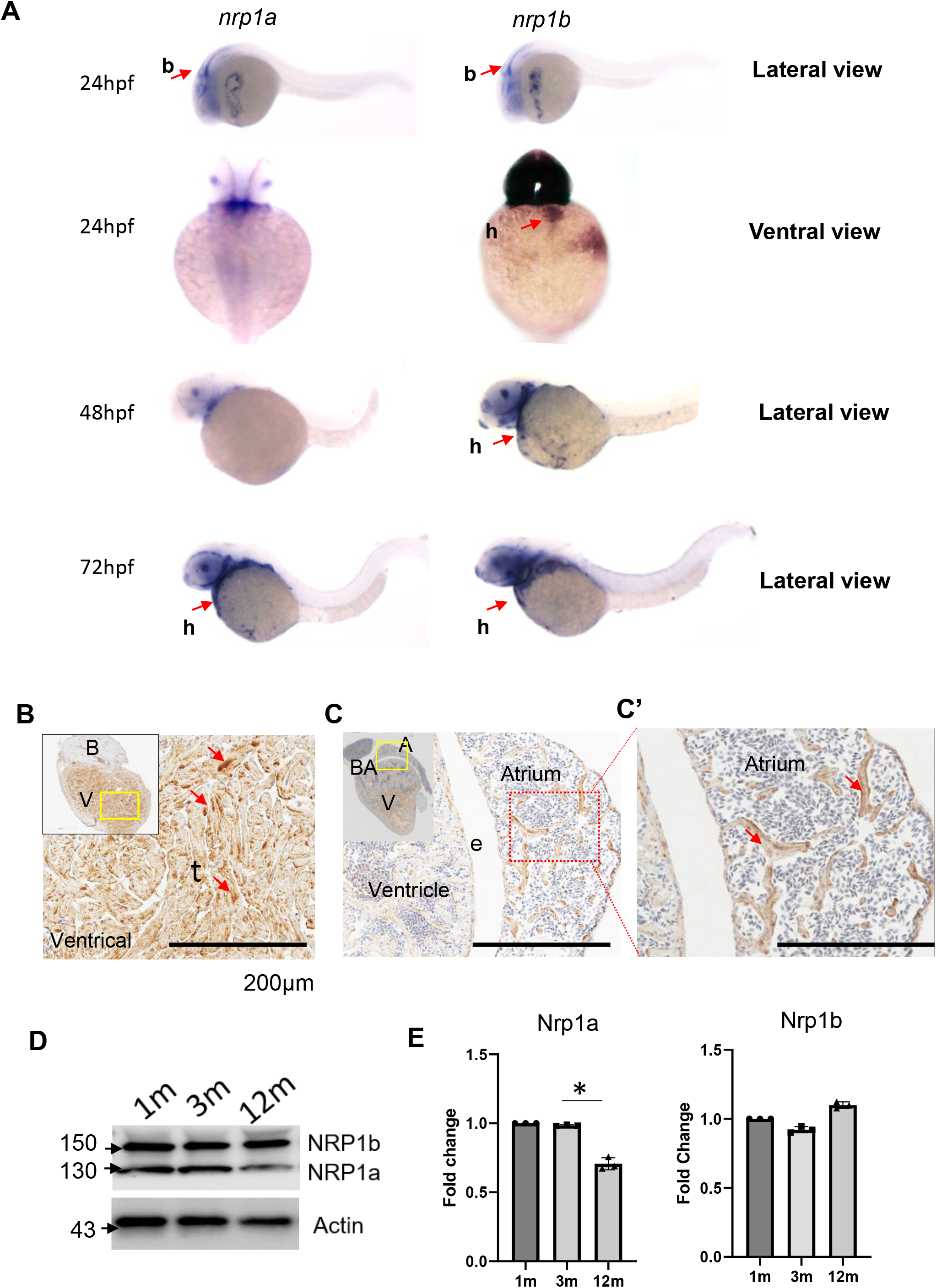
Nrp1 expression in the zebrafish embryo and the adult heart. **(A)**. Representative *In-situ* hybridization image showing nrp1a and nrp1b expression in 1 dpf, 36 hpf, 2 dpf and 3 dpf zebrafish embryo (Red arrowheads indicate the expression in the heart. (**B, C,** and **C’)** Immunohistochemistry image showing NRP1 expression in adult zebrafish heart (Yellow and the red dotted box indicates the area under focus in the enlarged image in B, C, and C’, and the red arrowhead indicates the NRP1 expression as indicated by the staining), (**D**) Western blot showing the NRP1a and NRP1b expression in adult heart at 1, 3 and 12 months (m). (E) Quantification of D. Error bars represents the mean ± standard deviation from three different experiments.*, p<0.05, **, p<0.01. Scale bar = 200μm. b, brain; h, heart; e, endothelium; t, trabecula. Statistical significance was evaluated with Prism 9.0 software by using a nonpaired, two-tailed Student’s t-test. P values below 0.05 were considered significant and, if lower, as indicated.

Further, Western blotting of the protein lysates prepared from 1-month, 3 months, and 12 months old zebrafish heart ventricles confirms that both the nrp1 isoforms are expressed in adult hearts. The Western blot showed two bands at ~130 kDa and 150 kDa, corresponding to Nrp1a [916 amino acids (aa)] and Nrp1b (959 aa), respectively (Fig 1D).

As shown in Fig 1E, the quantification of the Western bands showed a differential protein expression level of the Nrp1a isoforms at the three stages of development (1 mo, 3 mo, and 12 mo) in this study. The *nrp1b* isoform showed consistent expression levels in all three stages analyzed. In order to test the specificity of the NRP1 antibody used in our study, we have also performed the Western blot analysis to check the specific NRP1 knockdown in mouse CM treated with NRP1 shRNA. We tested two shRNA to target mouse cardiomyocytes (HL-1). As shown in Figure S1A, immunofluorescence staining confirmed the NRP1 knockdown in the cardiomyocytes after shRNA treatment. Western analysis also confirms the NRP1 downregulation after shRNA treatment (Fig S1 B-C). We also confirmed the cardiomyocyte specificity of this antibody in both adult zebrafish heart ventricles and mouse ventricle cardiomyocytes (HL-1) by immunostaining (Fig S2 and S3). Co-staining with cardiomyocyte marker MYH6 shows that NRP1 is expressed in the CM. In situ hybridization of each isoform in the knockout background also confirm the cardiomyocyte-specific expression of these isoforms. However, a detailed analysis will be required to confirm and verify the difference in the protein level at different stages of development, such as the early developmental patterns of these genes. It will be interesting to analyze the protein and mRNA expression levels of the isoforms at various developmental stages and the adult zebrafish heart. This expression profile will help explain these isoforms’ role in the corresponding stages.

### nrp1a and NRP1b are upregulated after cryoinjury in zebrafish adult heart

Next, we quantified how cryoinjury regulated the *nrp1a* and *nrp1b* mRNA levels in ventricles using qPCR. The *nrp1a* and *nrp1b* mRNA levels were upregulated on day 1, continued to increase 3-5 folds on day 3, and restored to normal levels at day 14 post cryoinjury (Fig. 2A and B). In situ hybridization analysis shows cardiomyocyte specific expression of both nrp1a (Fig S4 A, B) and nrp1b (Fig S4 C, D). Figure S4 E and F confirms an enriched expression of cardiomyocyte specific nrp1a and nrp1b levels after the injury, suggesting a cardiomyocyte specific role in zebrafish heart after cryoinjury.

**Figure 2.**
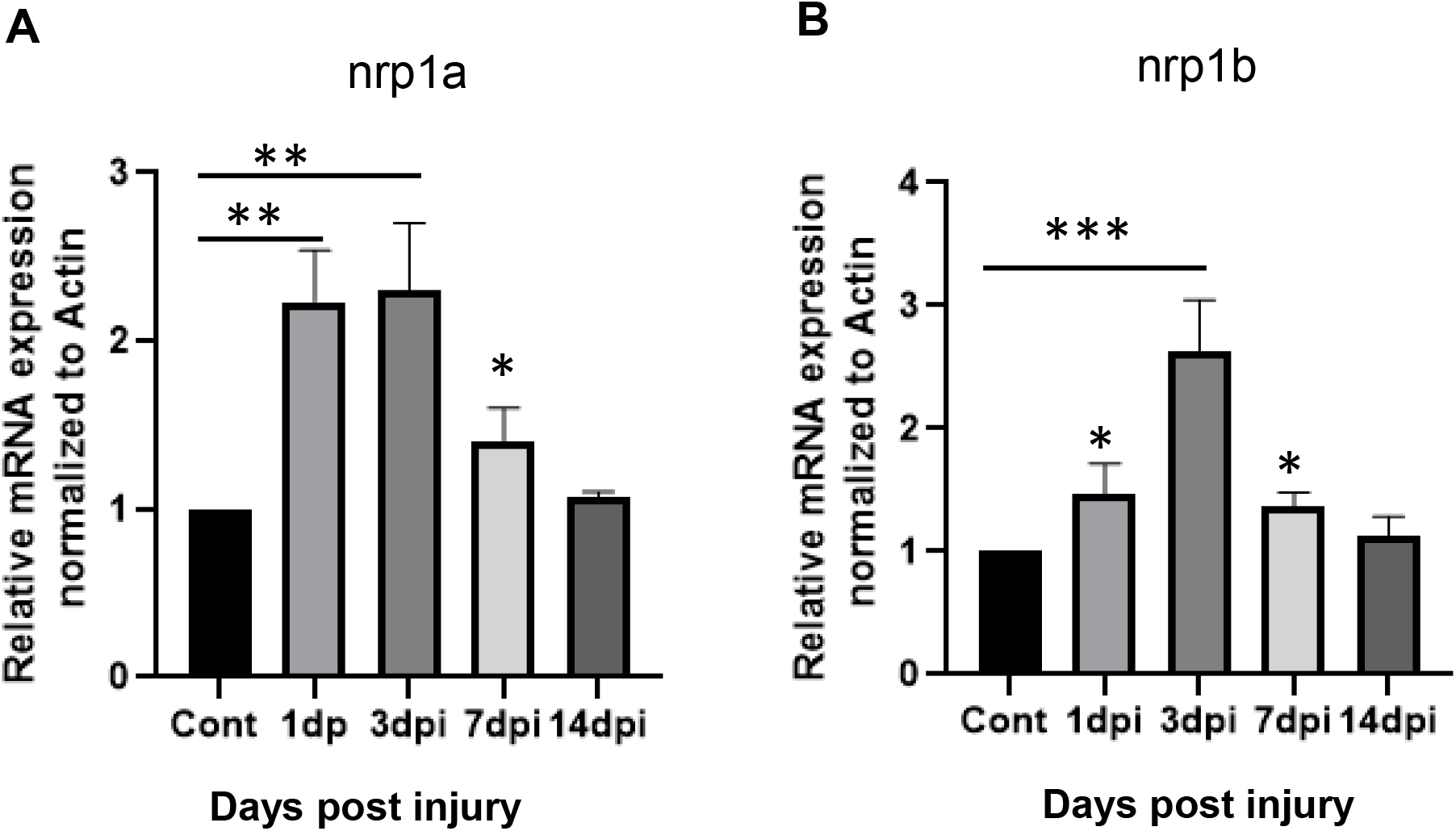
Differential expression of nrp1a and nrp1b in the injured heart. **(A).** RT PCR showing the upregulation of nrp1a mRNA expression after injury in zebrafish adult heart**. (B)** RT PCR showing the upregulation of nrp1b mRNA expression in adult zebrafish hearts after cryoinjury. Briefly, the RNA was isolated after 1d, 3d, 7d, and 14 d of cryoinjury from both the control and injured heart, and the relative mRNA expression was analyzed. The error bars represent the mean ± standard deviations. The experiments were repeated three times (biological replicate). * p<0.05, ** p<0.01 and ***, p<0.001. Statistical significance was evaluated with Prism 9.0 software by using a nonpaired, two-tailed Student’s t-test. P values below 0.05 were considered significant and lower as indicated.

### Construction of an integrative CRISPR vector for gene targeting in zebrafishfheat

To study the spatiotemporal role of Nrp1 isoforms in the zebrafish heart, we developed a conditional tissue-specific CRISPR Cas9 system. We developed a Tol2 integrative vector by modifying a previous vector developed by Ablain et al., 2015. As shown in Figure 3A: 1) a zebrafish heat shock promoter (Hsp70) drives the expression of a nrp1a or nrp1b gRNA scaffold, into which zebrafish codon-optimized Cas9 flanked by two nuclear localization signals are cloned^20^. The sgRNA is placed under the control of the Hsp70 promoter, which was cloned by using the Gateway cloning technology^21^, and green fluorescence protein (GFP) was identified downstream of the Cas9, which serves as a transgenesis marker^19^. Here cardiac myosin light chain (*cmlc2*) promoter drives the expression of the Cas9 flanked by EGFP. Using the *cmlc2* promoter helps in cardiomyocyte specific expression of Cas9 mRNA (Fig 1A).

**Figure 3.**
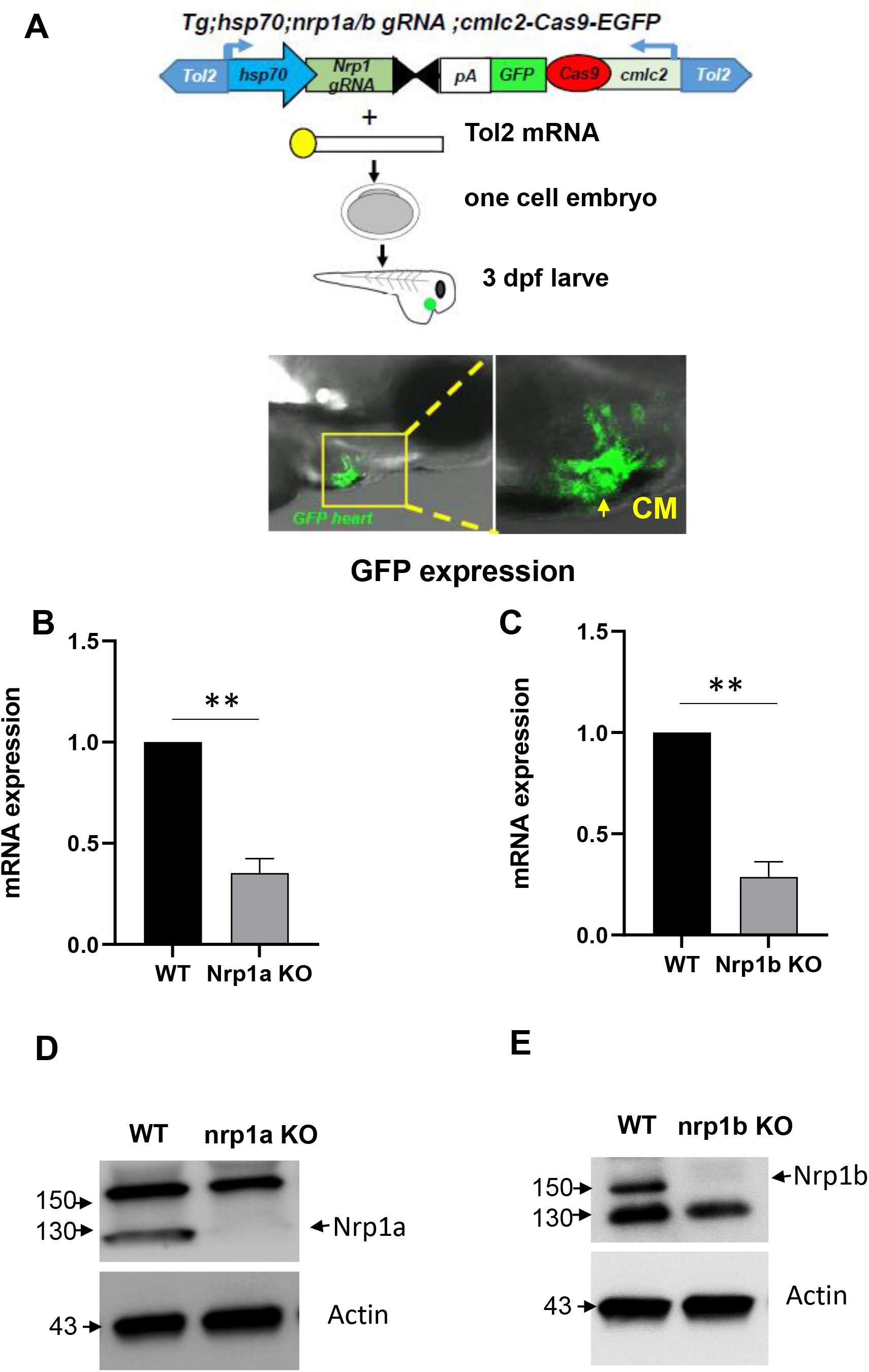
Outline of the Crispr-Cas9 mediated nrp1a and nrp1b knockout. (**A)**. Schematic showing *nrp1a* and *nrp1b* targeted Crispr-Cas9 construct, and tol2 mRNA mediated integration into the 1-2 cell zebrafish embryo confirmed by the green fluorescence protein expression in the heart. (**B and C**) Relative mRNA expression of *nrp1a* and *nrp1b* showing *nrp1a* and *nrp1b* knockdown after heat shock induction in adult zebrafish heart. (**D and E**) Western blotting showed nrp1a and nrp1b protein expression knockdown in the adult zebrafish heart at 3 months (3m). *Hsp70;* heat shock protein promoter, which drives nrp1 gRNA expression, and *cmlc2; Cardiomyocyte specific*, cardiac myosin light chain 2 promoters which drive the Cas9 expression in the heart. The green fluorescence protein expression in the heart confirms cas9 expression. *Tol2*; Tol2 transposase, SV40 polyA sequence (pA). The RT PCR and the western experiments were repeated at least three times (biological replicates), and the error bars indicate the mean ± standard deviation. ** p<0.01. Statistical significance was evaluated with Prism 9.0 software by using a nonpaired, two-tailed Student’s *t*-test. P values below 0.05 were considered significant and, if lower, as indicated.

AB zebrafish strain embryos were injected with vectors, along with Tol2 mRNA, at 1 cell stage. The fluorescent signal was observed specifically in the heart as early as 24 hpf, and it becomes more prominent at 3 dpf as shown in Fig 3A. We did not observe any toxicity upon expression of our vectors in embryos apart from that usually associated with the microinjection of DNA plasmids at the one-cell stage.

**Figure 4.**
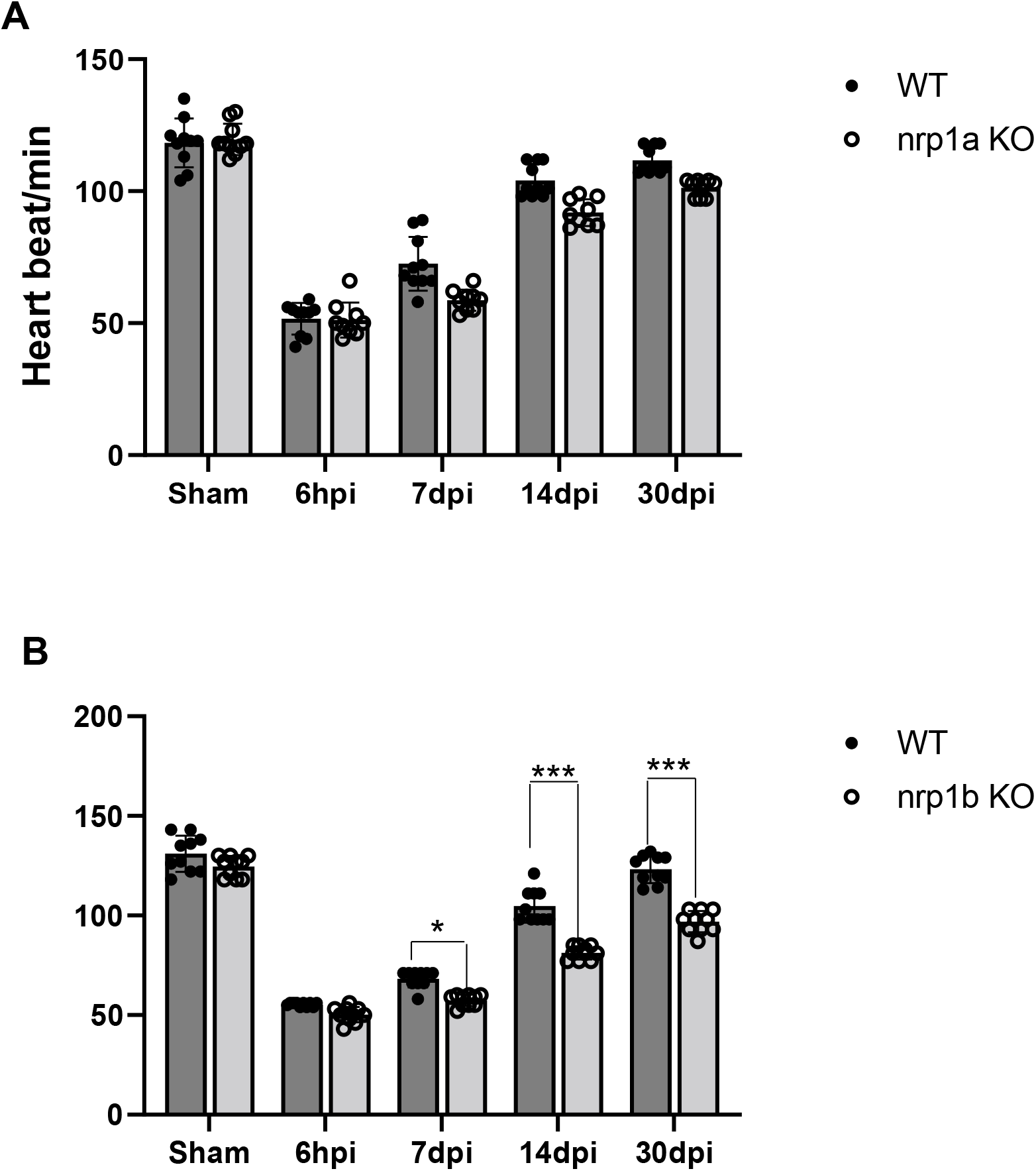
Heart rate analysis in 6-month-old WT and NRP1 KO zebrafish.x. **(A)**. Graph showing the heart rate recorded from n=10, WT, and Nrp1a KO fish. **B**. Graph showing the heart rate recorded from n=10, WT, and Nrp1b KO zebrafish embryo.. The bars represent the average values, and the error bars represent the mean ± standard deviation. * p<0.05, *** p<0.001. Statistical significance was evaluated with Prism 9.0 software by using Two-way ANOVA (multiple comparisons).

### The CRISPR vector allows tissue-specific gene disruption in zebrafish

To determine the efficiency of gene targeting, we first performed the T7E1 mutagenesis assay^29^. In whole embryos, significant mutation rates at the *nrp1a* and *nrp1b* target locus were only detected with the vector containing a gRNA against nrp1a or b and the Hsp70 promoter (Fig S5A-B). Injection of vectors expressing Cas9 under the control of the cardiomyocyte-specific *cmlc2* promoters and with the Hsp70 promoter driving the expression of a gRNA against *nrp1a* and *nrp1b* disrupts the nrp1 gene and protein expression after 72 hours post heat shock induction for 30 minutes as depicted by the mRNA expression (Fig 3 B and C), and protein level (Fig 3 D and E). Sanger sequencing confirmed the genome editing (Fig S5B). Briefly, we applied heat shock to zebrafish by placing them in 37°C water bath for 30 minutes as described in the method, followed by tissue collection 72 hours later. As shown in Fig S6A, nrp1a knockout zebrafish have reduced nrp1a expression in the heart. Similarly, nrp1b knockout zebrafish have reduced nrp1b levels in the heart at 2 dpf (Fig S6B). These data indicate successful ablation of the nrp1a and nrp1b genes in the adult zebrafish heart after heat shock.

### nrp1a and nrp1b abrogation have differential effects on zebrafish cardiac function after cryoinjury in the adult heart

To determine whether *nrp1a* and *nrp1b* abrogation alters heart function, we next performed a noninvasive electrocardiogram (ECG) to analyze the heart rate in the 10 −12 month-old zebrafish before and after cryoinjury. The mean R-R intervals were used to evaluate the heart rate variation. In the wild-type zebrafish, decreased heartbeats were observed at 6 hpi and slowly recovered to normal levels at 14 dpi (Fig 4 A-B). Deletion of *nrp1a* did not significantly affect the heartbeats before and after the cryoinjury (Fig. 4A). Although *nrp1b* deficient zebrafish showed a similar level of heartbeats at baseline and at 6 hpi and 7 dpi, their heart rates were significantly reduced at 14 dpi and 30 dpi, compared to WT zebrafish (Fig. 4B). The heat shock is known to affect the heart rate. However, in the current study, In the current study, we applied heat shock for 30 min to induce the NRP1 gRNA expression, and the functional analysis was carried out after 3 days when zebrafish are likely to recover from heat shock.

Next, we analyzed the cardiac function by echocardiography. As shown in Fig 5 (A-B), we observed that in the WT zebrafish, both ejection fraction (EF) and fractional shortening (FS) were reduced by 48% ± 2.2% at baseline to 26% ± 1.98% at 3 dpi, and then recovered to normal levels at 30 dpi indicating successful cardiac regeneration post-injury. Abrogation of the nrp1a isoform in the zebrafish heart does not affect the cardiac function post-injury, as shown by the similar level of EF and FS, compared with WT zebrafish. However, *nrp1b* deficient zebrafish showed delayed recovery of cardiac function and significantly reduced EF and FS at 30 dpi, compared to WT group (Fig 5 C-D). Our results suggested that cardiomyocyte specific-nrp1b but not nrp1a is required for the full recovery after cryoinjury in zebrafish.

**Figure 5.**
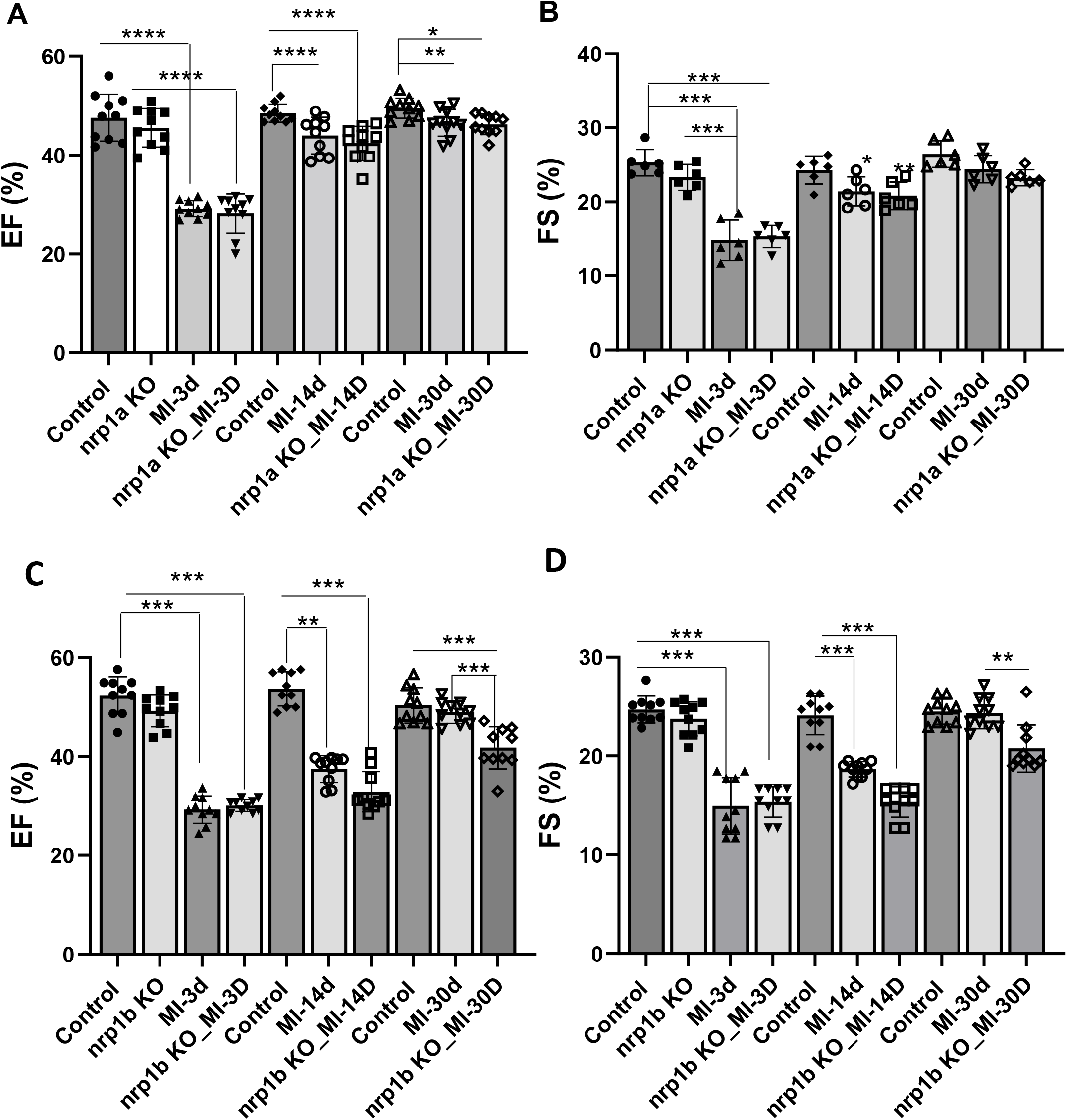
Heart function analysis in adult WT and nrp1a and nrp1b KO zebrafish. **(A)**. Graph showing the ejection fraction, recorded from n=10, WT, and Nrp1a KO fish. and (**B)**. Graph showing the fraction shortening recorded from n=06, WT, and Nrp1b KO zebrafish embryo. CRISPR-Cas9 mediated nrp1b knockout delayed the cardiac function recovery after injury in adult zebrafish heart. **(C)**. Graph showing the ejection fraction (EF), recorded from n=10, WT, and nrp1a KO fish. and (**D)**. Graph showing the fractional shortening (FS) recorded from n=10, WT, and nrp1b KO zebrafish embryo.. The bars represent the average value, and the error bars indicate the mean ± standard deviation. ** p<0.01 from three repeats. * p<0.05, ** p<0.01, ***, p<0.001 and ****p<0.0001. Statistical significance was evaluated with Prism 9.0 software by using a nonpaired, two-tailed Student’s t-test. P values below 0.05 were considered significant and, if lower, as indicated.

### nrp1b ablation results in enhanced expression of cardiac remodeling gene in adult zebrafish hearts after cryoinjury

Previously, we have shown that Nrp1 ablation in mouse cardiomyocytes upregulates the fetal gene expression (Wang et at., 2015)^30^. To investigate the effects of nrp1a, and nrp1b expression on fetal gene expression in 8 - 9 month-old zebrafish heart, we measured the cardiac hypertrophy and stress markers, including the brain natriuretic peptide (Bnp), atrial natriuretic factor (Anf/Nppa), Myosin heavy chain 6 (Myh6), and the Troponin gene (Tnnt2). As shown in Fig 6 A, the expression of Bnp, Nppa, and Myh6 were not changed upon nrp1a ablation in the nrp1a knockout group as compared to the WT, at both baselines and after cryoinjury. On the other hand, we observed that in the nrp1b knock-out zebrafish, the expression of Bnp, Myh6, and Nppa were upregulated at both baselines and after cryoinjury compared to the WT zebrafish (Fig 6 B). The expression level of the *tnnt2* was not changed at baseline but downregulated after cryoinjury in both the WT and nrp1b knockout zebrafish (Fig. 6B). These results suggest that nrp1 isoforms play a differential role in regulating the homeostasis of CMs after cryoinjury in zebrafish. These results also indicate that deletion of nrp1b aggravated the expression of cardiac remodeling genes after cryoinjury.

**Figure 6.**
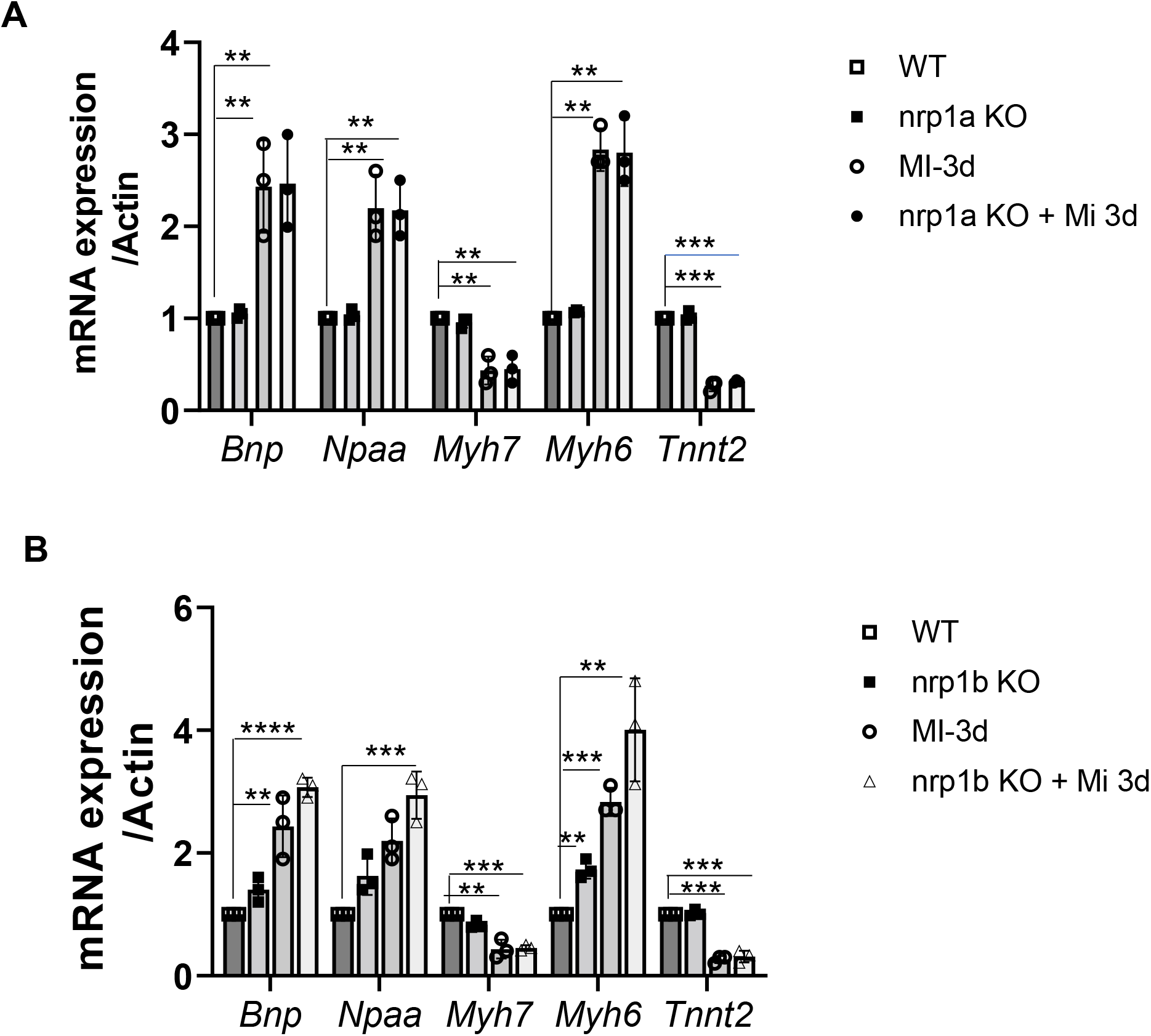
Nrp1a and Nrp1b abrogation results in differential expression of cardiac remodeling gene expression in adult zebrafish hearts. Nrp1a and b have a different regulatory effects on cardiac remodeling in adult zebrafish hearts. **(A)** mRNA expression of cardiac remodeling genes in heart isolated from the adult 8–9-month-old wt and nrp1a KO zebrafish after 3 days post cryoinjury(3dpi). (**B**). mRNA expression of cardiac remodeling genes in heart isolated from the adult 8–9-month-old wt and nrp1b KO zebrafish heart after 3 days post cryoinjury (3dpi). The RT PCR experiments were repeated three times (N=5 hearts were combined to isolate the RNA in each group), and the error bars indicate the mean ± standard deviation. ** p<0.01 *** p<0.001, **** p<0.0001. Statistical significance was evaluated with Prism 9.0 software by using a nonpaired, two-tailed Student’s t-test (N = number of experiments or animals analyzed). P values below 0.05 were considered significant and if lower as indicated.

### nrp1b ablation delays cardiac regeneration in zebrafish

To further confirm the effect of cardiomyocyte specific nrp1a and nrp1b on cardiac regeneration, 6-month-old WT, nrp1a, and nrp1b knockout zebrafish were subjected to cryoinjury, and heart ventricles were analyzed for scar area (the size of the trichome staining by histology at 3 dpci, 14dpci, and 30 dpci. (The percentage area of the collagen-rich scar was calculated relative to the area of the ventricle (Fig. 7). At 3 dpi and 14 dpi, deletion of nrp1a or nrp1b did not significantly affect the scar sizes. However, at 30 dpi, nrp1b deficient zebrafish exhibited significantly increased scar areas compared with WT group (5.2% ± 1.48% vs. 3.7% ± 0.98%), whereas nrp1a knockout zebrafish did not affect the scar area (p=0.038, Fig. 7A and B). These results indicate that cardiomyocyte specific nrp1b is required to resolve the scar and cardiac regeneration.

**Figure 7.**
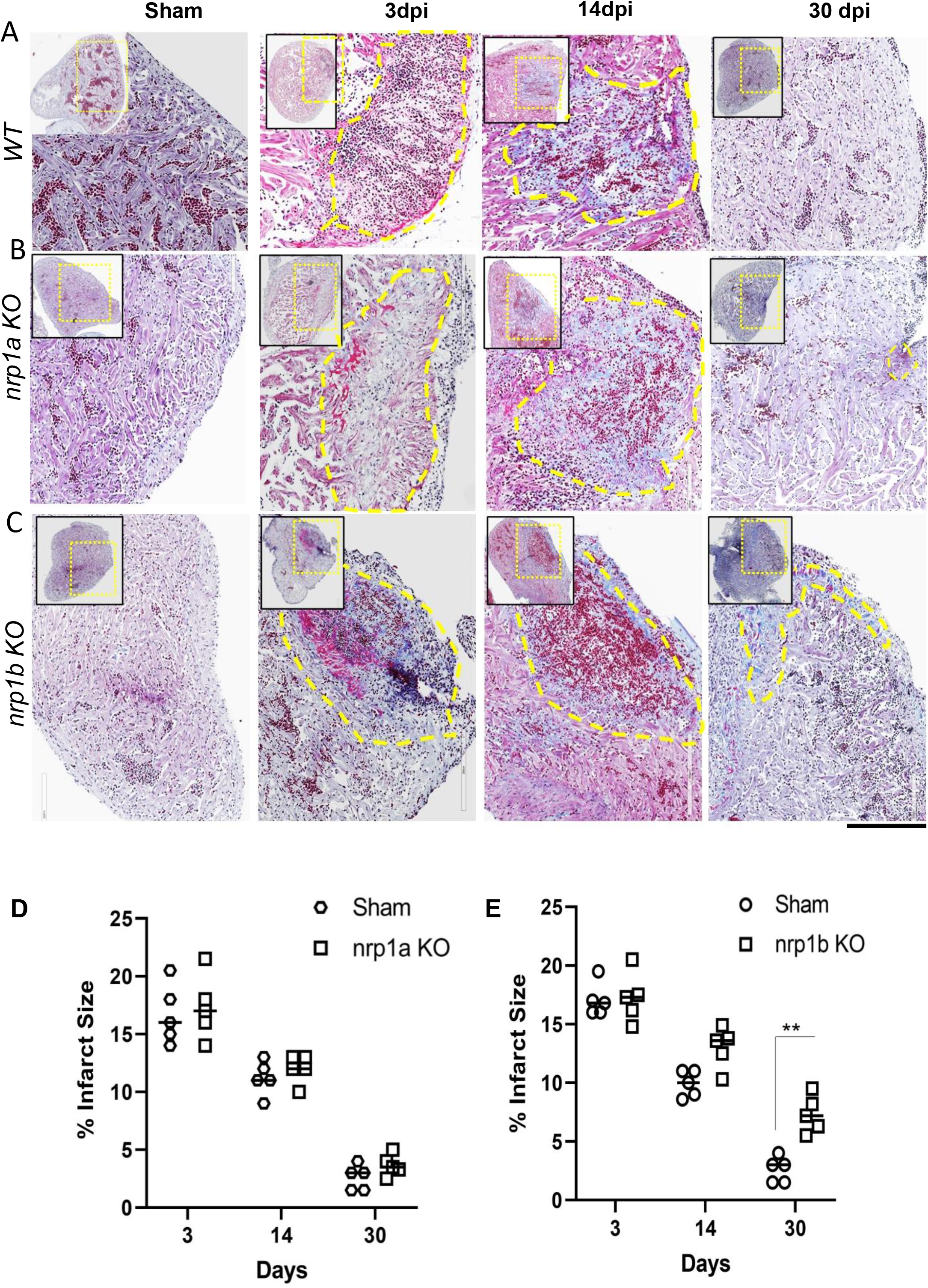
Analysis of the infarct size in nrp1a and nrp1b KO zebrafish after cryoinjury. **(A-C)** Representative heart cross-section of Sham, 3 dpi, 14 dpi, and 30 dpi heart from the top of the ventricle (left upper corner) to the ventricular apex (right bottom corner) labeled to trichome staining to visualize the healthy myocardium in purple, fibrin in red and collagen in blue in WT, nrp1a KO, and nrp1b KO zebrafish. (**D**) Quantification of the infarct area in WT vs nrp1a KO zebrafish. **(D)** Quantification of the infarct area in WT vs nrp1b KO zebrafish. The yellow dotted box indicates the region of the heart analyzed in the enlarged image, and the yellow dashed regions indicate the infarct area. **v**, ventricle; **ba**, bulbus arteriosus; **a**, atrium. The bars indicate the average values, and the error bars indicate the mean ± standard deviation. The quantifications were performed by measuring the infarct area from five different samples in each timepoint.. **, p<0.01. Statistical significance was evaluated with Prism 9.0 software by using a nonpaired, two-tailed Student’s *t*-test (N = number of experiments or animals analyzed). P values below 0.05 were considered significant and if lower as indicated.

### Nrp1b abrogation inhibits the proliferation of cardiomyocytes

BrdU proliferation assay and PCNA staining in the *nrp1a* and *nrp1b* knockout fish after cryoinjury was performed to investigate further the role of nrp1b depletion in the proliferation. Our result confirmed that *nrp1b* abrogation significantly reduced the number of proliferating cells as compared to the *nrp1a* and wild type fish heart. The result has confirmed that nrp1a and wild type zebrafish heart shows no significant difference in proliferation (Figure S7). We also check the effect of nrp1a and nrp1b ablation on neovascularization. As shown in Fig S8 A and B, nrp1a and nrp1b ablations reduced the number of new vessels formed in the injury site.

### nrp1b ablation abrogates the MMP9 gene expression in zebrafish heart

Previously, it has been shown that matrix metalloproteinases (MMPs) activity is critical during the inflammatory phase of heart regeneration^31^. To see whether nrp1 ablation alters the expression of MMPs, we measured the mRNA expression of mmp2, *mmp9* and mmp13 in the RNA samples isolated from the nrp1a and nrp1b knockout zebrafish hearts after three day post cryoinjury. While we did not observe significant changes in the expression of *mmp9* when *nrp1a* was downregulated (Fig 8A), there was an approximately two-fold decrease in the *mmp9* mRNA level in *nrp1b* isoform knockout zebrafish (Fig 8B). We observed that mmp2 but not mmp13 expression was reduced in the nrp1b knockout zebrafish heart (Fig S9 A). Additionally, we have tested the mRNA expression of various markers, including *vegfaa, vegfr2, vegfr1, vegfc, pdgfab, tgfb1*, and *vimentin* at early timepoint (1dpi). We observed increased expression of *vegfr2, vegfaa, vegfc, pdgfab* and *tgf1* after 1dpi in the wild type. We also show upregulation of *vimentin* expression at 1dpi in the wild type zebrafish. We evaluate the expression of these genes to study the role of angiogenesis and the expression of genes associated with epithelial-to-mesenchymal transition which are implicated in the regeneration process^16^. *Nrp1b* depletion was found to inhibit the upregulation of *pdgfab*, *vimentin*, and *tgfb1* after injury at 1dpi (Fig S9 B-E).

**Figure 8.**
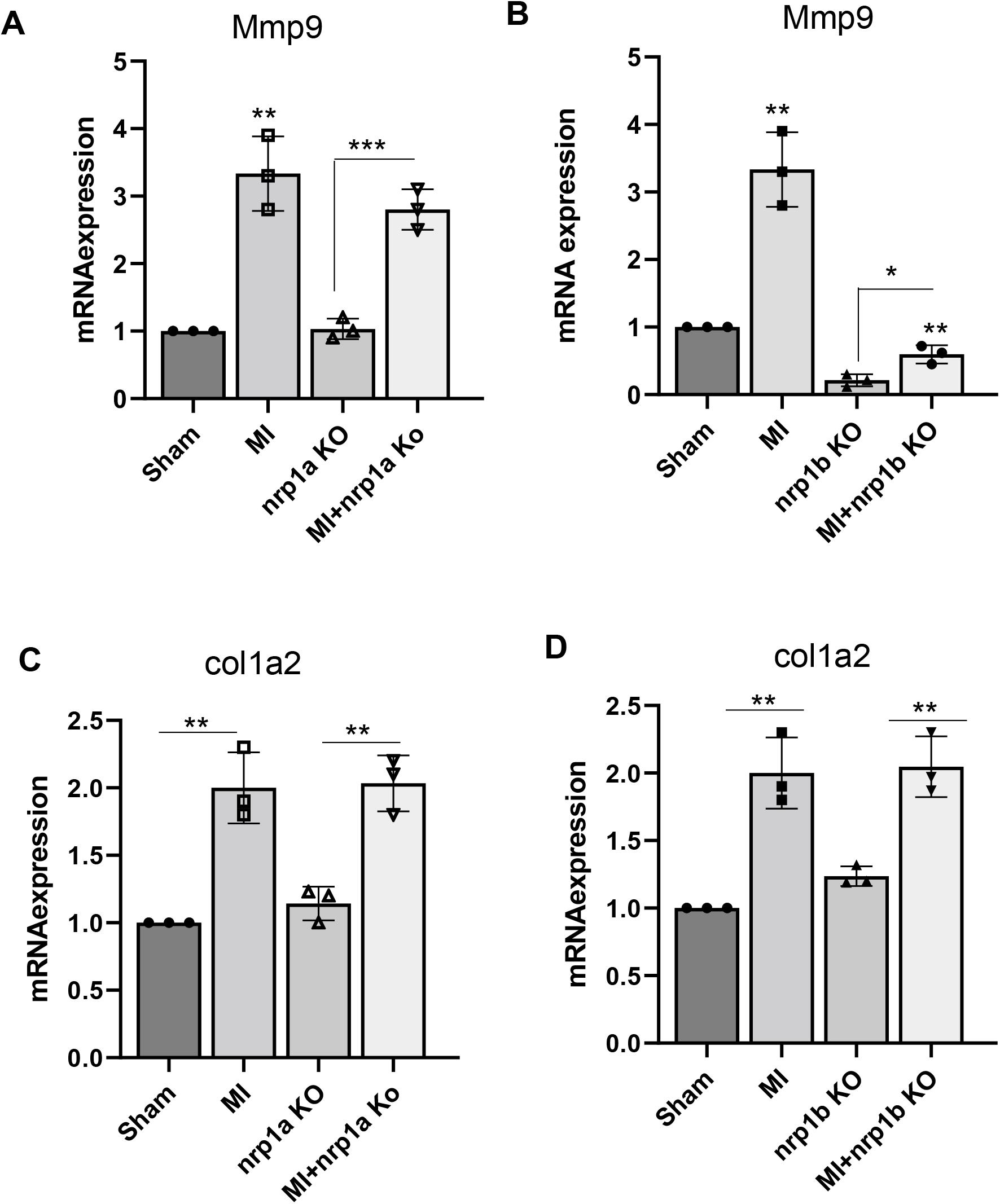
nrp1b abrogation inhibits the expression of mmp9 expression in adult zebrafish hearts. **(A-B)** Graph showing the relative mRNA expression of mmp9 in **(A)** WT and nrp1a KO. **(B)** WT and nrp1b KO zebrafish heart after cryoinjury. **(C-D).** Graph showing the relative mRNA expression of *col1a2* in **(C)** WT and nrp1a KO **(D)** WT and nrp1b KO zebrafish heart after cryoinjury. Error bars represent the mean ± standard deviation, and the RT PCR experiments were repeated three times. *, p<0.05, **, p<0.01, and ***, p<0.001. Statistical significance was evaluated with Prism 9.0 software by using a nonpaired, two-tailed Student’s *t*-test (N = number of experiments or animals analyzed). P values below 0.05 were considered significant and lower, as indicated.

To confirm the direct regulation of MMP9 by NRP1, we have transfected mouse ventricle cardiomyocyte (HL-1) cells with NRP1 shRNA and analyzed the mRNA expression of MMP9. We did observe reduced RNA expression of MMP9 in the NRP1 shRNA transfected cells (Fig S10A). Previous studies have suggested that regeneration-associated cardiac fibrosis influences the heart regeneration process. In this study, we measured the expression of the other ECM gene Col1a2 (collagen synthesis gene), a well-known fibrosis marker. We observed that *nrp1a* or *nrp1b* knockout did not significantly affect the expression of Col1a2 mRNA. These results indicate that MMP9 is involved in the delayed cardiac regeneration of *nrp1b* deficient zebrafish.

### Nrp1a and nrp1b isoforms doesn’t compensate each other

Analysis of the mRNA and protein level of each isoform when the other is abrogated. Doesn’t display significant changes in the expression of one isoform upon the removal of another isoform (Figure S10B). This result suggested no compensatory mechanism between the two isoforms. Importantly, when we abrogated the expression of *nrp1b* isoform, *tgfb1*, and *pdgfab* mRNA expression were also reduced. The same was true for PDGF expression when nrp1a was downregulated. However, *tgfb1* expression remains unchanged after nrp1a downregulation (Fig S9B-E). These results suggested a differential role of *nrp1a* and *nrp1b*.

## Discussion

To increase the potential applications of the effective CRISPR-Cas technology, we have developed a targeting vector that allows for tissue-specific inactivation of a gene in a spatiotemporal manner by modifying a previously developed vector^15^. We show that the CRISPR-Cas knockout technology can be spatially controlled in zebrafish and our vector system allows for the generation of stable zebrafish lines with heart-specific gene knockout. We have exploited this technology to overcome the challenges in the embryonic lethality associated with the global knockout of essential genes, which confounds the analysis of their functions in animal models. Moreover, since the approach is Tol2 transposon technology-based, our approach may apply to model organisms other than zebrafish.

Both the *nrp1a* and *nrp1b* mRNAs were expressed in the zebrafish heart with differential levels across the development, and both the isoforms were strongly upregulated in the zebrafish heart 1 −3 days after cryoinjury. These expressions were shown to coincide with the epicardial activation time, which occurs very early following cardiac injury^16^. It was observed that the upregulation of nrp1 isoforms was maintained for 3-7 days after the cryoinjury, further supporting the conclusion that nrp1 is involved in the early phase of regenerating heart post cryoinjury. Lowe et al. have suggested that these isoforms play an essential role in the epicardial activation phase, which occurs during the first 3-7 days of regeneration^16^ Our finding did observe an important role of nrp1a in early time point (3-14 dpi) of heart regeneration as reported by Lowe et al 2019. But the role of cardiac specific nrp1b isoform which has not been examined before in the process of heart regeneration in zebrafish hearts. We noticed that the cardiac-nrp1b isoform showed a more harmful effect on regeneration than the nrp1a isoform and wild type.

One major difference is that Lowe et al. used the global nrp1a deficient zebrafish model, whereas we used the heat inducible-cardiomyocyte-specific nrp1a knockout zebrafish, suggesting that the role of nrp1a in cardiac regeneration is spatiotemporal dependent. Given that impaired cardiac regeneration is only observed in the global but not cardiomyocyte-specific nrp1a knockout zebrafish model, it is likely that the non-cell autonomous role of nrp1a is required for cardiac regeneration. On the other hand, we have utilized a cardiomyocyte specific nrp1a and nrp1b knockout system, which restrict the nrp1 deletion in the cardiomyocyte in an inducible manner. Moreover, we did observe similar inhibitory effects of both nrp1a and nrp1b ablation in the cryoinjury led neovascularization (Figure S8) which was also reported by Lowe et al.^16^. Our results suggested nrp1b/MMP mediated regulation in the injured heart. Our study advocates the use of tissue specific ablation of nrp1a and nrp1b isoforms will help to understand their cell-specific function. For our study, we have focused on the specific role of nrp1a and nrp1b in cardiomyocytes in the overall heart function recovery after injury. Despite the advancement in the field, not many studies have described the role of nrp1a and nrp1b in cardiac repair after injury.

Analysis of CRISPR-Cas9 mediated *nrp1a*, and *nrp1b* knockout zebrafish provided direct evidence that nrp1b is required for zebrafish heart regeneration. As previously described^16^, the nrp1a knockout fish did not display any morphological defect or pathological phenotype. Similarly, we did not observe any lethal phenotype in both *nrp1a* and *nrp1b* adult knockout fish. It was previously reported that ablation of nrp1 using morpholino oligomers produces a lethal phenotype in zebrafish embryos (Martyn and Schulte-Merker, 2004). Lowe et al. and we have shown that the spatiotemporal knockdown of nrp1 in adult zebrafish didn’t cause lethality, allowing us to explore the functions of these genes in adult cardiac repair after injury. Following cardiac injury, the CRISPR mediated, cardiomyocyte specific *nrp1a* and *nrp1b* knockout fish demonstrated a differential effect on heart regeneration. The *nrp1b* knockout significantly lowered regenerative function compared to WT controls and the *nrp1a* knockout group. The importance of *nrp1b* for heart regeneration was demonstrated by the delayed and incomplete removal of fibrin deposits essential for the scar resolution process in the knockout zebrafish. The delayed wound closure in *nrp1b* fish probably indicates a failure of the myocardium to migrate efficiently towards the subepicardial layer and to prevent fibrosis after cryoinjury, as suggested by Lowe et al., 2015^16^. These findings support that *nrp1b* is required for zebrafish heart regeneration following cryoinjury. We have checked the mRNA expression of both isoforms in the *nrp1a* and *nrp1b* knockout zebrafish heart. We didn’t observe any compensatory role of one isoform on the other. It is possible that cardiomyocyte specific *nrp1a* and *nrp1b* loss mediated by double knockout would have an even more harmful role in the heart regeneration and function of zebrafish. Furthermore, our data show that *nrp1b* regulates the *mmp9* expression and thus indicating a possible role of *mmp9* in fibrosis during the repair process in heart regeneration. Cardiomyocyte specific NRP1 depletion in mice is accompanied by the upregulation of cardiac remodeling genes, including ANF, BNP βMHC^17^. Positive regulation of MMP9 by NRP1 in trophoblast cell proliferation and migration is known^32^. NRP1 ablation results in decreased protein and mRNA levels of MMP9^32^.

Moreover, the role of the myofibroblasts is essential in post-MI healing. MMP9 promotes cardiac fibroblast migration, increased collagen production, increases angiogenic factors, and stimulates the transition of cardiac fibroblasts to myofibroblasts^33^. ECM produced by myofibroblasts contributes to tissue replacement and scar formation after the MI. Importantly, MMP9 has been associated with cardiac remodeling, and targeted deletion of mmp9 in mice leads LV remodeling^34–36^. Mechanistically, mmp9 inhibition resulted in reduced enzymatic activity which is responsible for ECM breakdown. In zebrafish, mmp9 and mmp13 are shown to play key roles in the inflammatory phase of heart regeneration following cryoinjury^31^. A recent study from Xu et al. showed an increase in MMP enzymatic activity and elevated expression of MMP9 and MMP13 in the injured area of hearts from as early as 1 day post-cryoinjury (1 dpi)^31^. Inhibition of MMP9 resulted in impaired heart regeneration, as indicated by the larger scar and reduced numbers of proliferating cardiomyocytes (Shian Xu 2018)^31^. Together these results and our data suggested nrp1b regulation of MMP9 in the cardiac remodeling in regeneration post cryoinjury. These findings warrant further investigation into the nrp1/MMP axis in regulating cardiac regeneration post-injury.

## Supporting information

Supplemental matterial

## Acknowledgment

The authors would like to thank Dr. Leonard Zon for providing the CRISPR vector clones. We thank Dr. Pritam Das for reviewing the manuscript and providing his helpful suggestions. We thank Dr. Laura lewis Twiffin for help in Confocal imaging. We thank Brandy Enfiled for her help in immunostaining. We thank Dr. Tanmay Kulkarny for his help in Raman microscopy.

## Author contribution

DM developed the concept and interpreted the data, and acquired funding. RA designed the experiment, analyzed the data, and drafted the manuscript. YW designed the experiments, revised the manuscript, and acquired funding. RA, SD, and EW performed the experiments.

