## Supplemental matterial for "Conditional, tissue-specific CRISPR/Cas9 vector system in zebrafish reveal the role of neuropilin-1b in heart regeneration"

Debabrata Mukhopadhyay, PhD

Department of Biochemistry and Molecular Biology

Mayo Clinic

4500 San Pablo Road South

Jacksonville

FL 32224

Figure S1

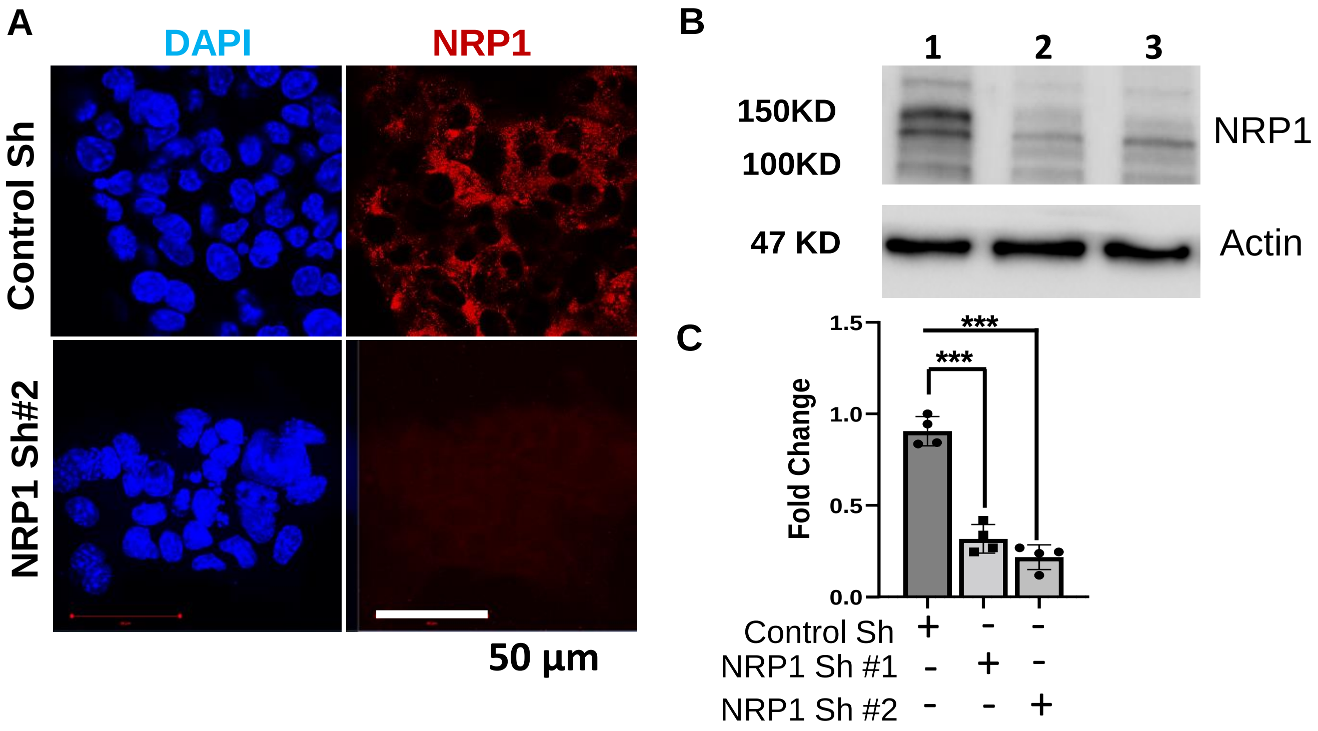

**Figure S1: NRP1 antibody validation in Cardiomyocytes. (A).** Immunofluorescence staining of Control shRNA and NRP1 shRNA treated mouse cardiomyocytes (HL-1) with NRP1 antibody (red) (Neuropilin-1 (D62C6) Rabbit mAb #3725). (**B**). Western blotting showing the NRP1 downregulation in HL-1 cardiomyocyte infected with NRP1 shRNA #1 and NRP1shRNA #2. (**C**). Quantification showing the fold change in protein expression from B. Error bars represents the mean ± standard deviation. The western experiment was repeated three times. ***, $p<0.001$. Statistical significance was evaluated with Prism 9.0 software by using nonpaired, two-tailed Student's *t*-test. P values below 0.05 were considered significant and if lower as indicated.

Figure S2

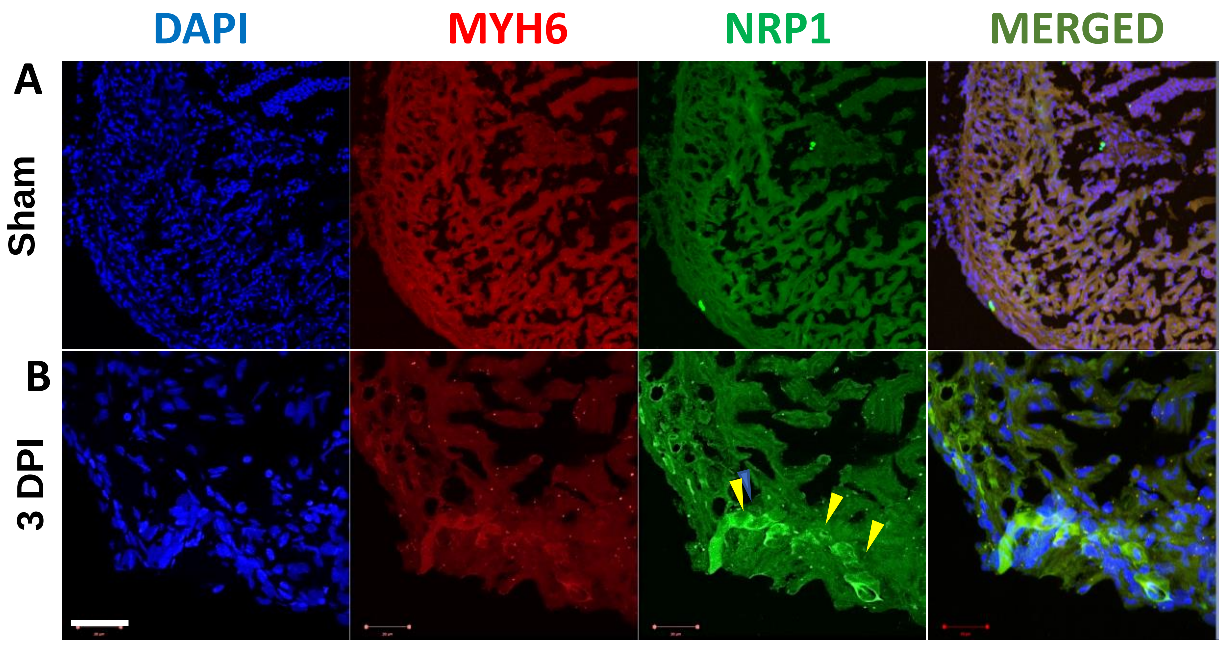

**Figure S2: NRP1 is upregulated in the injured cardiomyocytes**. **(A)** Confocal image showing NRP1 and MYH6 costaining in uninjured zebrafish ventricle cryosection and (**B)** Confocal image showing NRP1 and MYH6 costaining in zebrafish ventricle after injury (3 day after cryoinjury)**.** Yellow arrowhead indicated the NRP1 over-expression at injury site. Scale = 20µm

Figure S3

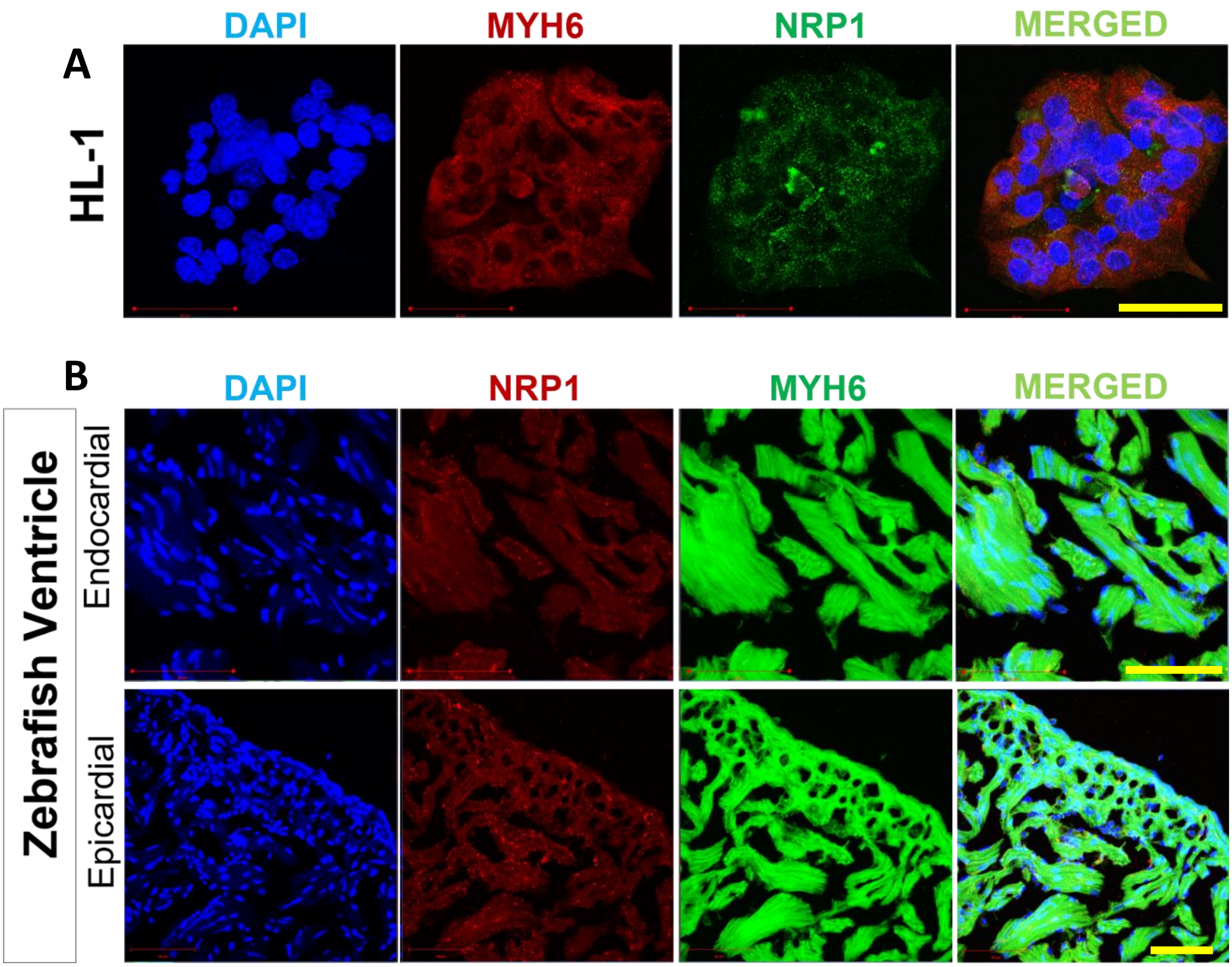

**Figure S3: NRP1 expression in cardiomyocytes.** (**A**). Confocal image showing NRP1 and cardiomyocyte marker, MYH6 (Anti-MYH6 antibody [3-48] ab15) costaining in mice ventricle cardiomyocyte (HL-1). (**B**). NRP1 and cardiomyocyte marker (MYH6) costaining in adult zebrafish ventricle. Scale = 50µm.

Figure S4

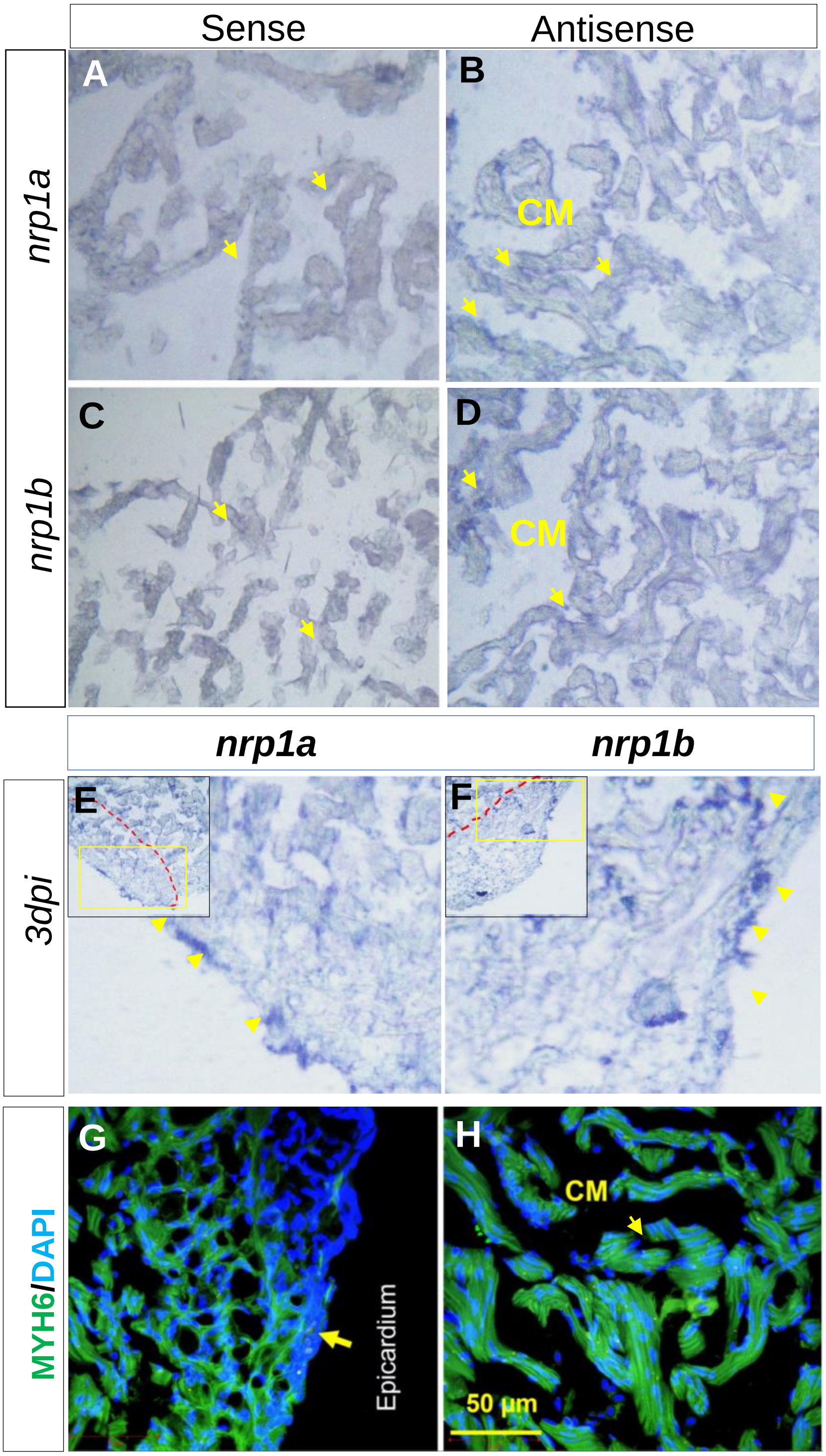

**Figure S4: Cardiomyocytes specific expression of nrp1a and nrp1b in zebrafish. (A-B).** In situ hybridization showing nrp1a expression in adult wild type heart ventricle section (A; control, B; target probe). (**C-D)**. nrp1b expression in adult wildtype heart ventricle (C; control, D; target probe). (**E).** In situ hybridization showing nrp1a expression in heart ventricle section after 3 dpi. (**F)**. nrp1b expression in heart ventricle section after 3 dpi. (**G-H)**. Confocal image of zebrafish ventricle cryosection stained with cardiomyocytes-specific myosin heavy chain 6 (green) and nuclear-DAPI. ( G; Periphery and H; inner region). Image were captured by Renishaw, Raman Microscope in A - F, Zeiss confocal microscope in G and H. Yellow arrowhead indicates the cardiomyocyte. Yellow arrow indicates epicardium. Yellow box indicate the enlarged area analyzed. Scale bar = 50µm, N= 5 images were analyzed for each group.

Figure S5

**A**

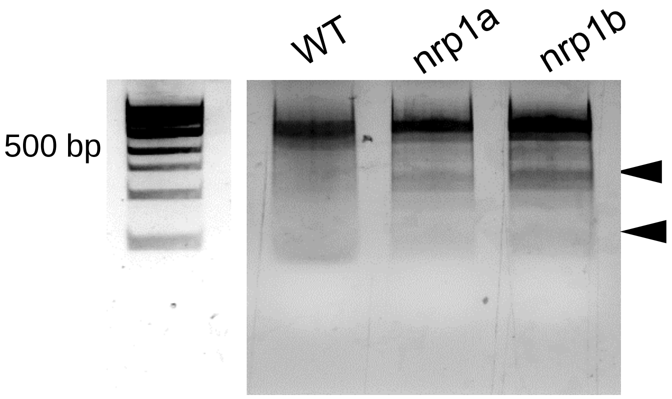

**B**

*nrp1a*

ACCCGATACCTGCGGATCCGACCCATTAACT**GGGAAACTGGCATCGCACTG**CGGTTCGAGGTCTACGGATGCAAG – WT

CTGACCCGATACCTGCGGATCCGACCCATTAACT**GGGAA**-----------**CACTG**CGGTTCGAGGTCTACGGATG

CTGACCCGATACCTGCGGATCCGACCCATTAACT**GGGAAA**------**TCGCACTG**CGGTTCGAGGTCTACGGATGC

CTGACCCGATACCTGCGGATCCGACCCATTAACT**GGGAAACTG**-----------------------------

*nrp1b*

TCCTAGTAGCATACTAGCGCTAGTGACTGTAGCTGCTGTA**GTCTCCATAGTAGTGGCAAG**AGGAAGAGG -WT

GTAGCATACTAGCGCTAGTGACTGTAGCTGCTGTA**GTCTCCA**-------**GGCAAG**AGGAAGAGGTGTTG

GTAGCATACTAGCGCTAGTGACTGTAGCTGCTGTA**GTCT**-------**GTGGCAA**GAGGAAGAGGTGTTGTG

GTAGCATACTAGCGCTAGTGACTGTAGCTGCTGTA**GTCT**-----------**GCAA**GAGGAAGAGGTGTTGT

GTAGCATACTAGCGCTAGTGACTGTAGCTGCTGTA**GTCT**-------------------------------------

**Figure S5: CRISPR cas9 mediated genome editing in nrp1a and nrp1b. (A).** T7E1 mutagenesis assay at the CRISPR target site in the nrp1a and nrp1b gene. The assay was performed on genomic DNA from 4 dpf embryos injected at the one-cell stage with Cas9 mRNA and either a gRNA against nrp1a or a gRNA against nrp1b. Cleavage bands (arrowheads) indicate the presence of mutations at the target site. (**B)**. Representative sequencing result showing the mutation sites.

Figure S6

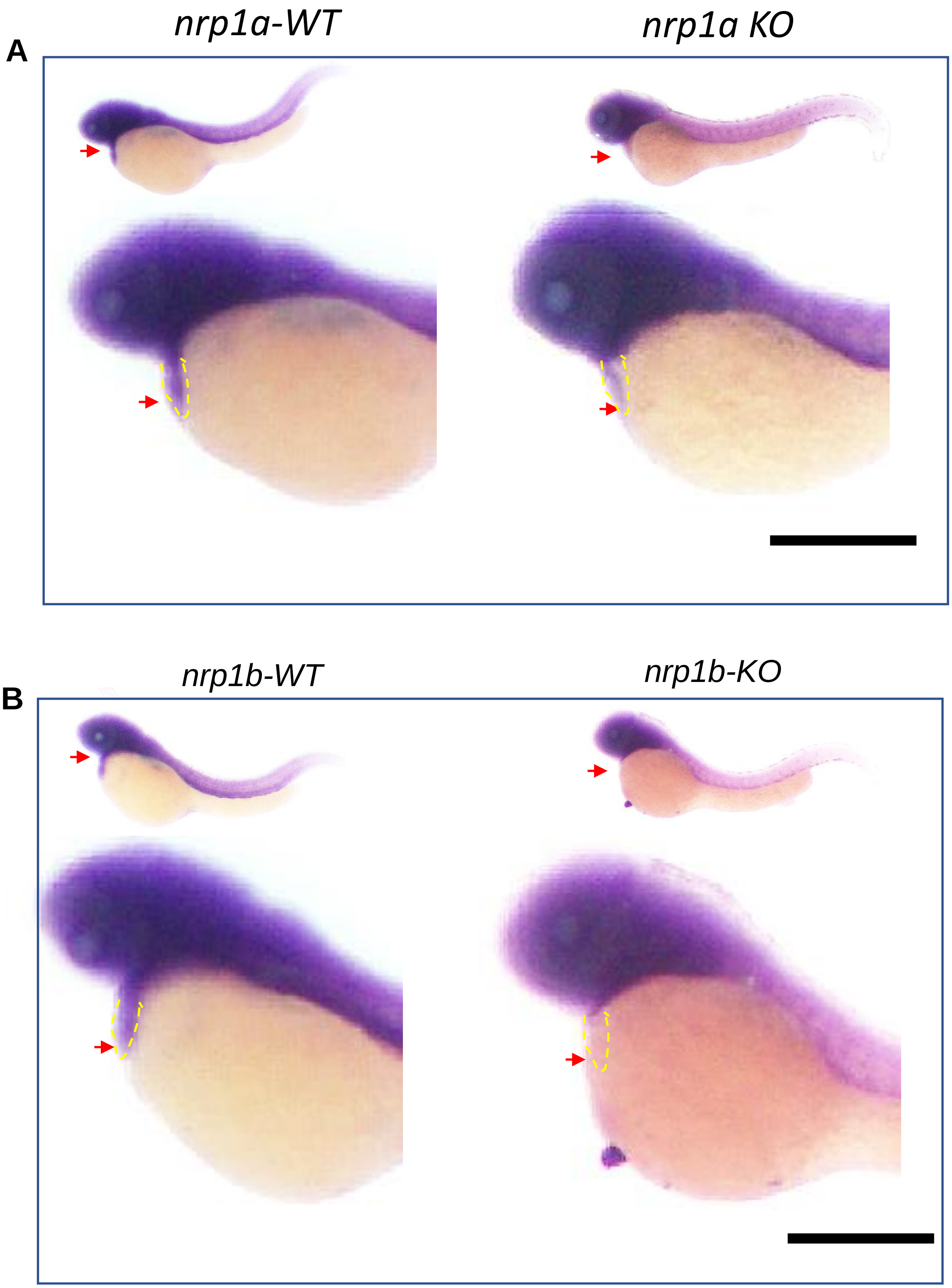

**Figure S6: Cardiac specific nrp1a and nrp1b deletion in 3 dpf zebrafish embryo**. **(A)** Representative image showing in situ hybridization with antisense nrp1a riboprobe in WT and nrp1 KO, 2 dpf zebrafish embryo. **(B)**. Representative image showing in situ hybridization with antisense nrp1b riboprobe in WT and nrp1b KO, 2 dpf embryo. Scale = 0.5mm. N=10 zebrafish embryos were analyzed in each group. Yellow dotted line indicates the heart specific expression of nrp1a and nrp1b.

Figure S7

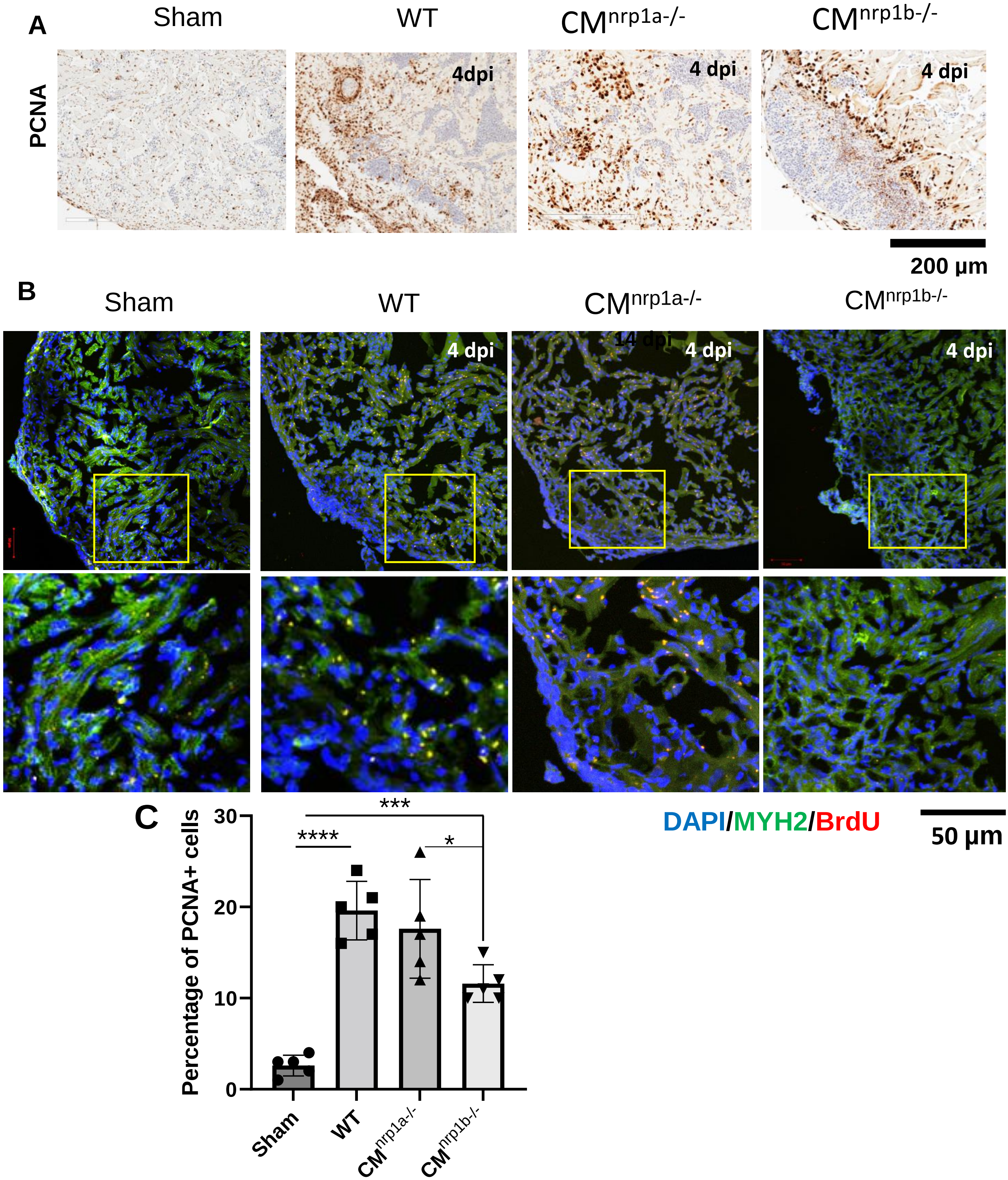

**Figure S7: nrp1b ablation reduced the proliferation of cardiomyocyte after cryoinjury. (A).** PCNA staining showing proliferative cells in WT-sham, WT- 4 dpi and CM specific nrp1a/nrp1b KO zebrafish adult heart section after 4 dpi. **(B)**. Representative confocal image showing the BrdU staining in WT- sham, WT-4 dpi and CM specific nrp1a/nrp1b KO zebrafish adult heart section after 4 dpi. **(C).** Quantification of the BrdU staining in B. Yellow square box indicates the analyzed area. Error bars represents the mean ± standard deviation (N=5 different images were analyzed for the quantification). ***, p<0.001. Statistical significance was evaluated with Prism 9.0 software by using nonpaired, two-tailed Student's *t*-test. P values below 0.05 were considered significant and if lower as indicated.

Figure S8

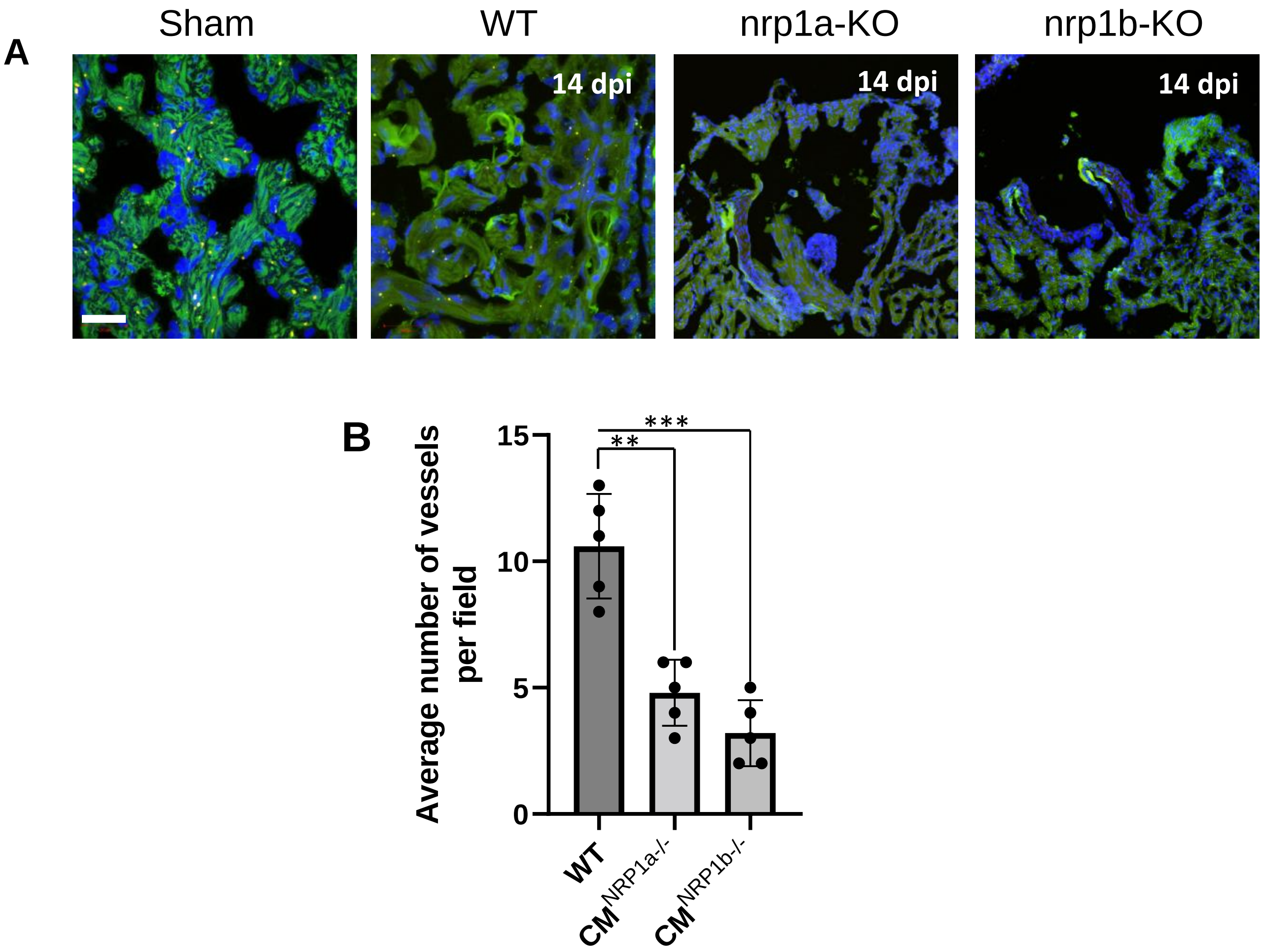

**Figure S8: Neovascularization of the cryoinjured area is impaired in cardiomyocyte specific *nrp1a and nrp1b KO*. (A).** Confocal image of ventricle section showing the *VE*-cadherin (green) in the WT-sham, WT- 4 dpi, and CM specific nrp1a/nrp1b KO zebrafish adult heart section (4 dpi). (**B)**. Quantification showing the number of neo-vessels in the four groups. Scale bar= 20µm. Error bars represents the mean ± standard deviation (N=5 images were analyzed per group). **, p<0.01, and ***, p<0.001. Statistical significance was evaluated with Prism 9.0 software by using nonpaired, two-tailed Student's *t*-test. P values below 0.05 were considered significant and if lower as indicated.

Figure S9

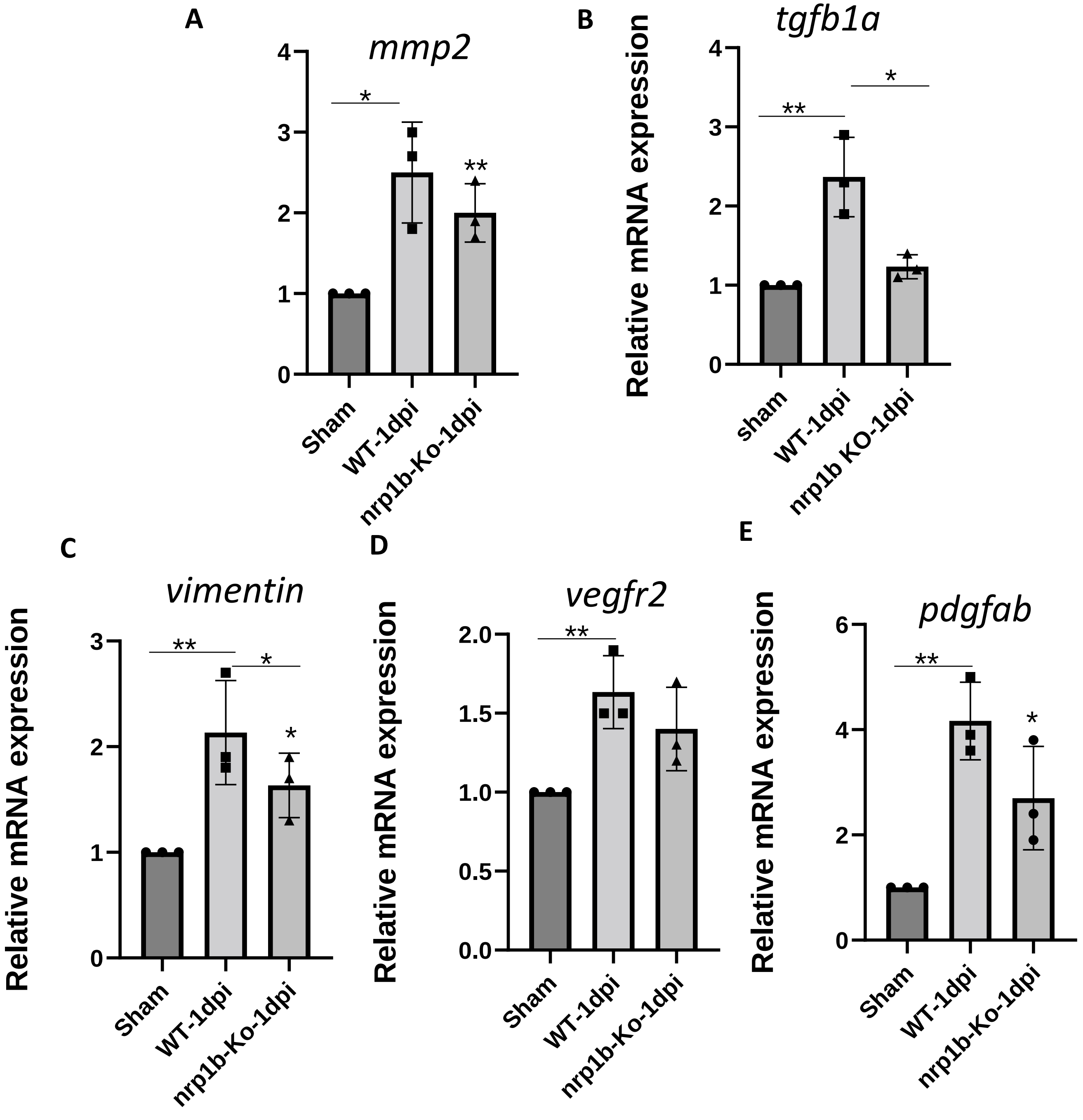

**Figure S9: RNA expression of MMP2 and vimentin in the injured heart. (A)** Relative mRNA expression of MMP2 in control, cardiac specific nrp1a and nrp1b KO heart after cryoinjury (1 dpi). (**B and C)** Relative mRNA expression of EMT marker genes expression in control, cardiac specific nrp1a and nrp1b KO heart. **(D and E)** Angiogenesis marker in. Error bars represents the mean ± standard deviation and the experiments were repeated at least three times. *, p<0.05, **, p<0.01. Statistical significance was evaluated with Prism 9.0 software by using nonpaired, two-tailed Student's *t*-test. P values below 0.05 were considered significant and if lower as indicated.

.

Figure S10

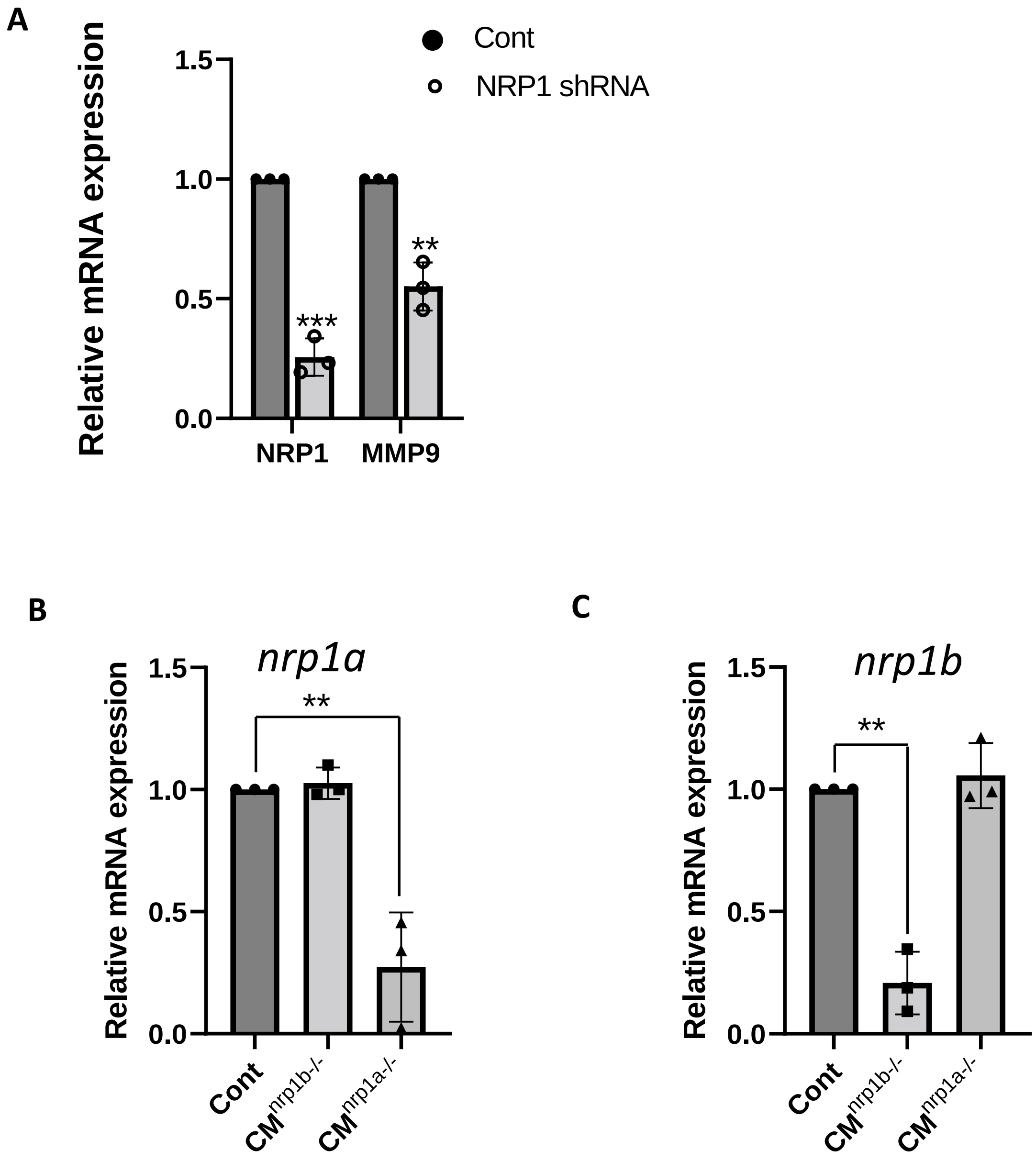

**Figure S10: Compensatory effect *nrp1a and nrp1b*. (A)** Relative mRNA expression of Nrp1 and MMP9 in control, and NRP1 shRNA#2 transfected mice ventricle cardiomyocyte (HL-1). **(B).** Relative mRNA expression of nrp1a in control, cardiac specific nrp1a and nrp1b KO heart. **(C)**. Relative mRNA expression of nrp1b in control, cardiac specific nrp1a and nrp1b KO heart. Error bars represents the mean ± standard deviation and the experiments were repeated at least three times. **, p<0.01, and ***, p<0.001. Statistical significance was evaluated with Prism 9.0 software by using nonpaired, two-tailed Student's *t*-test. P values below 0.05 were considered significant and if lower as indicated.

.

### Supplemental Table 1. Primer sequences

| Gene | Forwards | Reverse |
|---|---|---|
| Mmp9 | TGATGTGCTTGGACCACGTAA | ACAGGAGCACCTTGCCTTTTC |
| Col1a2 | AAGAACCCCGCTCGTACTTG | TCCAGTAGAAACCGCTGCTC |
| Mef2ca | CCGTCCATGAACATGAGCCT | ACCGGCTCCGACTTAATGTG |
| Mef2cb | GTACAACGAGCCACACGAGA | CACCTGCACTAGGTGGTCTG |
| Col1a1 | GGCTTCCAGTTCGAGTATGG | ATGCAATGCTGTTCTTGCAG |
| Anp | GATGTACAAGCGCACACGTT | TCTGATGCCTCTTCTGTTGC |
| Vmhcl | TGTTGCAATCCAGACCGTCA | GCCACTTGTAGGGGTTGACA |
| Myh6 | GCATTCATTTCGGGACGAGC | R GACGTGAAGCCAAGCACATC |
| Tnnt2a | AGCAGAGCAGCAGAGAATCC | R AGAGGTTTGCGTCGATCACC |
| Myh7 | ATCAGGAGGTGGTTGTAGCC | GCAGGGTTAGCCTGGATGATT |
| nppa | GATGTACAAGCGCACACGTT | TCTGATGCCTCTTCTGTTGC |
| nppb | CATGGGTGTTTTAAAGTTTCTCC | CTTCAATATTTGCCGCCTTTAC |
| tnnt2c | GACCGAACGTGAGAAGAAGA | AGGACTTCCTGGTGGTTTTC |
| Actin | GCCTACTGGCCAGACGTCACAAATC | TCCAGCAAAACCGGCTTTGCACATAC |
| mouse MMP-9 | CTTCTGGCGTGTGAGTTTCCA | ACTGCACGGTTGAAGCAAAGA |
| MMP13 | CCTCCATATGAGGGCGTTGG | GATACATGAGTGCACCGGGA |
| pdgfab | CTGCTGCAACACCGGAAAC | GATCCTCTAACCGGACCAGC |
| tgfb1 | CAACCGCTGGCTCTCATTTG | CCTCTCTGCTTGTCTAGCCC |
| tnfa | AGACACGACCACAGCACTTC | CGGCACATTGCCAAGAGTGT |
| MMP13 | CCTCCATATGAGGGCGTTGG | GATACATGAGTGCACCGGGA |

### sgRNA sequence

| | |
|---|---|
| nrp1a | 1. AGAAATCCAAGATCAACCTGAGG<br>2. GGGAAACTGGCATCGCACTGCGG |

| | |
|---|---|
| nrp1b | 1. GTCTCCATAGTAGTGGCAAGAGG<br>2. GTCTCACACTAACCTTGGTGGTAGG |
